# Arc1 and the microbiota together modulate growth and metabolic traits in *Drosophila*

**DOI:** 10.1101/2020.07.21.213835

**Authors:** Scott A. Keith, Cassandra Bishop, Samantha Fallacaro, Brooke M. McCartney

## Abstract

Perturbations to animal-associated microbial communities (the microbiota) have deleterious effects on various aspects of host fitness, but the molecular processes underlying these impacts are poorly understood. Here we identify a novel connection between the microbiota and the neuronal factor Arc1 that affects growth and metabolism in *Drosophila*. We find that *Arc1* exhibits tissue-specific microbiota-dependent expression changes, and that germ-free flies bearing a null mutation of *Arc1* exhibit delayed and stunted larval growth, along with a variety of molecular, cellular, and organismal traits indicative of metabolic dysregulation. Remarkably, we show that the majority of these phenotypes can be fully suppressed by mono-association with a single *Acetobacter sp.* isolate, through mechanisms involving both bacterial diet modification and live bacteria. Additionally, we provide evidence that Arc1 function in key neuroendocrine cells of the larval brain modulates growth and metabolic homeostasis under germ-free conditions. Our results reveal a novel role for Arc1 in modulating physiological responses to the microbial environment, and highlight how host-microbe interactions can profoundly impact the phenotypic consequences of genetic mutations in an animal host.

**SUMMARY:** *Drosophila* Arc1 exhibits microbiota-dependent, tissue-specific differential expression and functionally interacts with a key *Acetobacter sp.* isolate to regulate developmental growth and metabolic traits.

## INTRODUCTION

The physiology and life history traits of animals are shaped in remarkable ways by interactions with commensal and beneficial microorganisms (the microbiota). For many metazoans, microbial symbionts play integral roles in post-embryonic development and physiology to yield fit and fertile adults (McFall-Ngai et al., 2013; Robertson et al., 2019). Thus, perturbations to the microbiota can profoundly disrupt these processes. For example, germ-free (GF; microbiologically sterile) or antibiotic-treated mice exhibit decreased body fat (Smith et al., 2007), abnormal intestinal epithelial architecture (Hayes et al., 2018), and neurodevelopmental defects (Sampson and Mazmanian, 2015). In humans, gut bacterial dysbiosis has been implicated in the pathogenesis of Type 2 diabetes (Larsen et al., 2010), obesity (Shen et al., 2013), autism (Gilbert et al., 2013), and other disorders. However, the molecular factors that actuate microbial influence on host physiology and development are incompletely understood. *Drosophila melanogaster* and its gut microbiota are an ideal model to discover such factors, given *Drosophila*’s extensive genetic resources and the technical ease of generating GF and gnotobiotic flies (Broderick and Lemaitre, 2012; Douglas, 2018).

From a screen to identify microbiota-responsive genes, we discovered altered levels of *Drosophila Activity-regulated cytoskeleton associated protein 1* (*Arc1*) transcript in GF flies. *Arc1* is a *Drosophila* homolog of mammalian *Arc/Arg3.1*, a master regulator of synaptic plasticity in the brain (Carmichael and Henley, 2018; Shepherd and Bear, 2011). *Arc* transcription is highly upregulated while the brain is encoding novel information into neural circuits (Chen et al., 2020; Guzowski et al., 1999). Accordingly, reduced *Arc* expression impairs learning ability in rodents (Guzowski et al., 2000; Shandilya and Gautam, 2020), and defects in human Arc have been linked to neurological and neurodevelopmental disorders, including Alzheimer’s disease (Bi et al., 2018), autism spectrum disorders (Alhowikan, 2016), and schizophrenia (Fromer et al., 2014). Both mammalian *Arc* and fly *Arc1* encode retroviral group-specific antigen-like amino acid sequences, and are predicted to have independently derived from ancient Ty3/Gypsy retrotransposons (Ashley et al., 2018; Campillos et al., 2006; Cottee et al., 2019; Pastuzyn et al., 2018). Recently, it was shown that Arc/Arc1 can self-assemble into capsid-like structures that package and transport mRNAs into cultured neuronal cell lines and across synapses *in vivo*, constituting a novel mechanism of cell-cell communication (Ashley et al., 2018; Erlendsson et al., 2019; Pastuzyn et al., 2018).

As with mammalian *Arc*, synaptic activity leads to strong transcriptional upregulation of fly *Arc1* (Guan et al., 2005; Mattaliano et al., 2007; Montana and Littleton, 2006; Mosher et al., 2015). At the larval neuromuscular junction (NMJ), synapse maturation and plasticity require Arc1 protein capsid-mediated transfer of *Arc1* mRNA from motoneuronal boutons to post-synaptic myocytes (Ashley et al., 2018). Further, *Arc1* loss-of-function mutants differentially express a repertoire of enzymes involved in central carbon metabolism, have altered metabolomic profiles, elevated fat levels, and are starvation resistant (Mattaliano et al., 2007; Mosher et al., 2015), suggesting that Arc1 controls metabolic homeostasis via unknown mechanisms.

In *Drosophila,* energy metabolism impacts developmental timing and whole-organism growth that occurs exclusively during the larval stages (Edgar, 2006). Developmental growth is diet-dependent and genetically regulated by multiple nutrient-responsive, inter-organ signaling systems, including (but not limited to) the insect-specific steroid hormone 20-hydroxyecdysone (20E; Buhler et al., 2018; McBrayer et al., 2007) and the functionally conserved insulin/insulin-like growth factor (IIS) pathway (Brogiolo et al., 2001; Rulifson et al., 2002; Edgar, 2006; Gilbert, 2008). As in mammals, the *Drosophila* microbiota plays a crucial role in dietary influence on metabolic and developmental processes. Certain taxa of commensal bacteria (*Lactobacillus* and *Acetobacter spp.*) promote metabolic homeostasis and growth through mechanisms including nutritional provisioning and promoting host signaling pathways like IIS (Chaston et al., 2014; Consuegra et al., 2020; Kamareddine et al., 2018; Keebaugh et al., 2018; Matos et al., 2017; Sannino et al., 2018; Shin et al., 2011). Interestingly, the microbiota can also reduce the severity of nutritional and developmental phenotypes resulting from host genetic mutations (Dobson et al., 2015; Ma et al., 2019). While connections between the microbiota, host metabolism, and diet are widespread, an understanding of the molecular mechanisms that drive these connections is lacking.

Here we uncover an interaction among Arc1, the microbiota, and host diet that impacts growth and metabolic traits in *Drosophila*. We show that *Arc1* transcript and protein levels change in tissue-specific patterns in GF flies, and loss of *Arc1* dramatically exacerbates the developmental growth delay of GF larvae. Further, we found that a single *Acetobacter* isolate is sufficient to restore normal larval development and other hallmarks of metabolic health to *Arc1* mutants, partly through a mechanism that involves conditioning the larval diet. Selective Arc1 expression in growth-regulating neuroendocrine cells suppresses the metabolic and developmental defects of GF *Arc1* mutants. Lastly, we provide evidence of both microbiota-dependent and -independent IIS and 20E dysregulation in *Arc1* null larvae. Together our data reveal an experimental system wherein a single microbiota member supports the health of a metabolically destabilized host, and demonstrates a previously unknown role for Arc1 in mediating the animal’s response to its microbial environment.

## RESULTS

### *Arc1* transcript and protein are sensitive to microbial condition

To identify microbiota-responsive neuronal factors, we previously conducted a transcriptomic screen to identify *Drosophila* genes that are differentially expressed in adult heads upon elimination of the microbiota. *Arc1* was among the genes most responsive to host microbial condition (Keith et al., 2019, preprint). Specifically, *Arc1* transcripts are elevated in the heads of adult wild-type *Drosophila* grown under germ-free conditions compared to flies grown in gnotobiotic (GNO) poly-association with a four-species bacterial community comprising two *Acetobacter* (*Acetobacter sp.*, *A. pasteurianus*) and two *Lactobacillus* (*L. brevis*, *L. plantarum*) isolates (Fig. 1A). *Arc1* was previously identified in published RNA-seq comparisons of microbiota-associated *vs.* GF adult guts (Bost et al., 2017; Dobson et al., 2016; Guo et al., 2014; Petkau et al., 2017). Consistent with these reports, and in contrast to the adult head, we found *Arc1* transcripts are decreased in the adult gut following microbiota removal (Fig. 1A). These microbiota-dependent transcript changes occur in multiple wild-type lines, but both changes were not observed in every line (Fig. 1A).

**Fig. 1.**
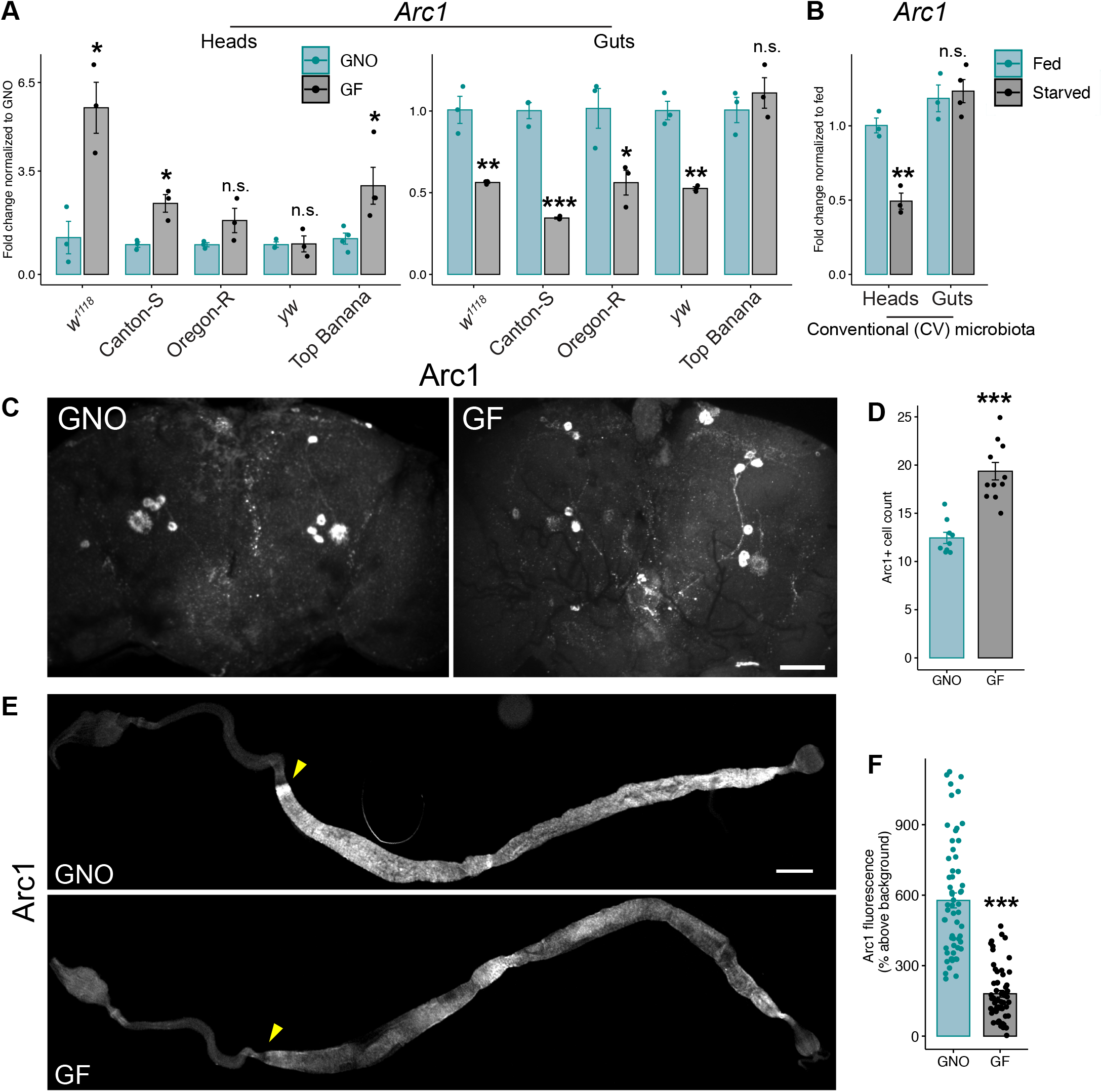
Tissue-specific *Arc1* mRNA and protein responses to microbiota and nutrient deprivation. Five-day-old adult male *w^1118^* flies were used for all analyses unless otherwise indicated. **A,B.** RT-qPCR of *Arc1* transcripts in heads and guts of adults grown GNO or GF (**A**), and following 24hr starvation (**B**). Individual points represent normalized values for each replicate, 10 animals per replicate, n=3-4. *p<0.05, **p<0.01, ***p<0.0001, n.s.=not significant, Student’s t-test. **C,D.** Arc1 immunostaining (**C**) and quantification of the number of Arc1-positive cells (**D**) in adult posterior central brains under GF and GNO conditions. Scale bar: 10 μm. Points represent cell counts from individual animals, n=9-11. ***p<0.0001, Student’s t-test. **E.** Arc1 immunostaining in the gut of GF and GNO flies, arrowheads indicate the midgut-hindgut boundary. Scale bar: 200 μm. **F.** Arc1 fluorescence intensity above background in posterior midgut cells at the midgut-hindgut boundary. Each point represents an individual cell, with 10 cells per animal, n=6 animals per condition. ***p<0.0001, Mann-Whitney test. Error bars=s.e.m.

Differential transcript abundances between microbiota-associated and GF animals could be a primary consequence of microbial loss, or may be secondary to metabolic changes resulting from sterile rearing. To test this, we measured *Arc1* levels in heads and guts of adult flies following nutrient deprivation, which alters metabolic function (Wat et al., 2020; Zinke et al., 2002). Interestingly, *Arc1* mRNA decreased ∼2 fold in the heads of flies starved for 24 hours, but was unchanged in starved guts (Fig. 1B), suggesting *Arc1* expression may be sensitive to nutritional inputs as well as the microbiota.

Prompted by our transcript-level findings, we examined Arc1 protein in the adult brain and gut under GNO and GF conditions. Our analysis of the brain revealed a complex, cell-type specific effect of microbial condition. On the posterior brain surface, there is a significant increase in Arc1-positive cells (Fig. 1C,D). While the level of Arc1 in the posterior central brain neurons was unaffected, Arc1 levels decreased in GF animals in a neighboring cell type with distinct morphology (Fig. S1A,A’’,C,D). By contrast, on the anterior brain surface, antennal lobe Arc1 expression increased in GF animals (Fig. S1B,E). *Arc1* transcript is reported to be highly enriched in the adult midgut (FlyAtlas; Leader et al., 2018), but its immunostaining pattern has not been reported. We found that Arc1 was broadly expressed throughout the foregut and midgut in GNO animals, with greatly reduced expression in the hindgut (Fig. 1E). Consistent with *Arc1* mRNA (Fig. 1A), Arc1 protein in midgut cells decreased ∼2 fold in GF animals (Fig. 1F).

Together these results show that Arc1 transcript and protein levels change in complex, tissue- and cell type-specific ways in response to microbiota and nutritional deprivation.

### Loss of the microbiota slows larval growth in *Arc1* mutants and a single *Acetobacter* species is sufficient to restore normal growth rate

*Arc1* regulates lipid homeostasis and central carbon metabolite levels in *Drosophila* larvae (Mosher et al., 2015), and loss of *Arc1* results in enhanced adult starvation resistance (Mattaliano et al., 2007). These and other metabolic traits are also impacted by the microbiota, depending on dietary conditions and host genetic background (Dobson et al., 2015; Gnainsky et al., 2021; Judd et al., 2018; Yamauchi et al., 2020). One major consequence of GF-induced metabolic dysregulation in *Drosophila* is prolonged larval growth (Strigini and Leulier, 2016). To test for an interaction between the microbiota and *Arc1*, we raised wild-type flies (*w^1118^*) and an *Arc1* deletion mutant (*Arc1^E8^*; Ashley et al., 2018) from embryo to adulthood either GF or GNO, and monitored larval growth rate. Consistent with many previous reports (reviewed in Strigini and Leulier, 2016), GNO wild-type animals completed larval development significantly faster than their GF siblings (∼7.3 days vs ∼8.8 days respectively; Fig. 2A, Table S1). Strikingly, the GF larval growth delay was dramatically extended in larvae lacking *Arc1*. While GNO *Arc1^E8^* mutant larvae developed at a rate indistinguishable from GNO wild-type, GF *Arc1^E8^* animals took on average ∼12.8 days to pupariate (Fig. 2A). These differences in larval growth rate were reflected in the rate of adult emergence (Fig. S2A). We observed a similar, though less protracted, developmental delay for two independently-generated *Arc1* loss-of-function alleles (*Arc1^esm18^* and *Arc1^esm113^*; Mattaliano et al., 2007) reared GF (Fig. S2B,C), and in GF *Arc1^E8^/Arc1^esm113^* transheterozygotes (Fig. 2B). All larvae bearing deletions of *Arc1* developed at the wild-type rate when grown with the GNO bacterial community. *Arc2* is another *Drosophila* Arc homolog in a genomic locus adjacent to *Arc1*; the two proteins likely represent an ancestral duplication event (Pastuzyn et al., 2018). In contrast to *Arc1* depletion, a P-element insertion in the *Arc2* 3’ UTR, which decreased *Arc2* expression (Fig. S2D), had no effect on the developmental rate of GF larvae (Fig. S2E).

**Fig. 2.**
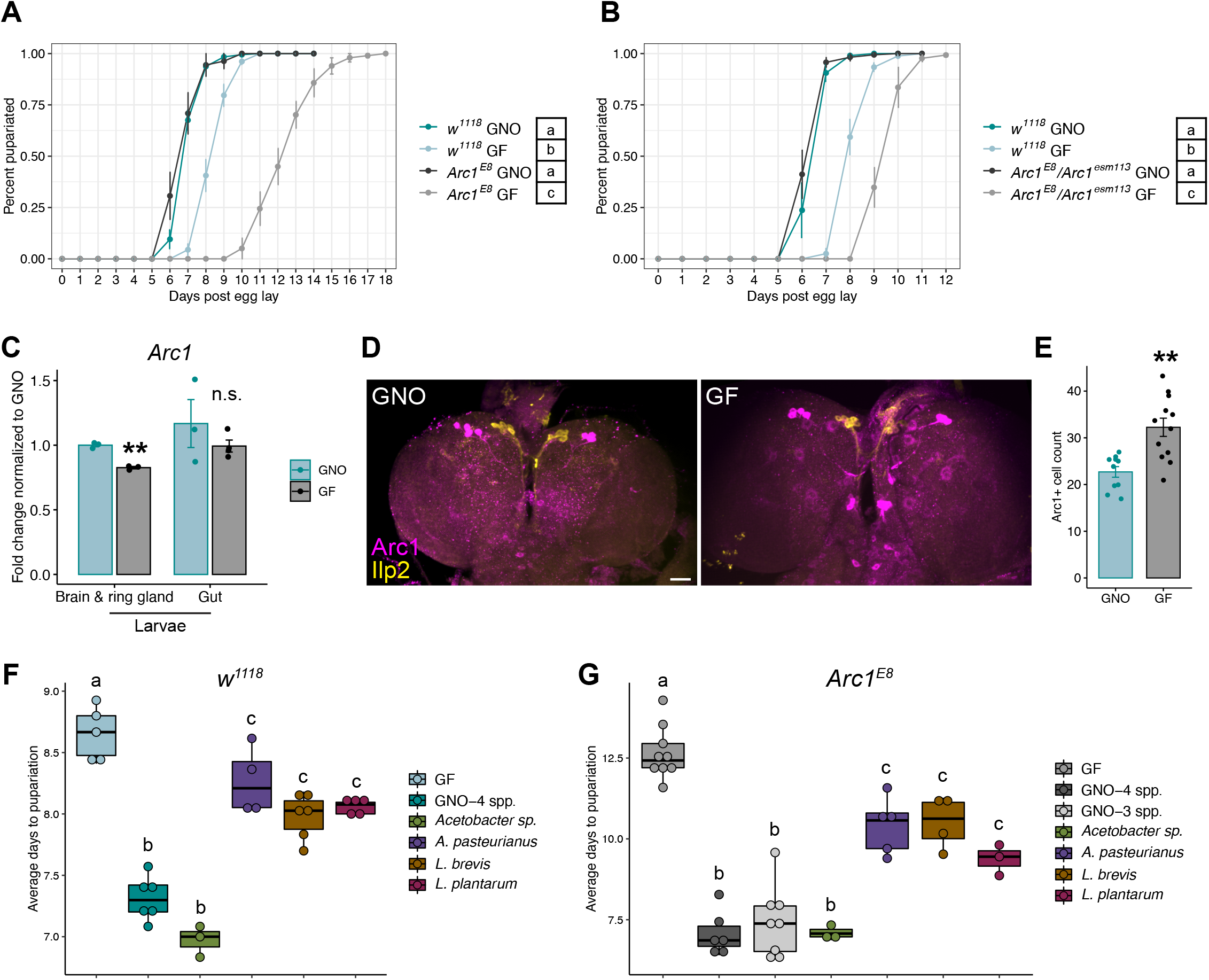
Mono-association with *Acetobacter sp.* promotes growth rate of *Arc1* mutant larvae. **A,B.** Time to pupariation for GNO and GF *w^1118^ vs. Arc1* mutants. **C.** RT-qPCR analysis of *Arc1* transcripts in indicated larval tissues. Individual points represent normalized values for each replicate, 10-20 animals per replicate, n=3-4. **D,E.** Arc1 and Ilp2 immunostaining **(D)** and counts of Arc1-positive cells **(E)** in the central region of the larval brain. Scale bar: 5μm. Points represent cell counts from individual animals, n=10-12. **F,G.** Time to pupariation for wild-type and *Arc1* mutant animals mono-associated with each of the four bacterial isolates comprising the GNO-4spp. condition. GNO–3spp.: *A. pasteurianus*, *L. brevis*, and *L. plantarum.* **(C,E)** **p<0.01, n.s.=not significant, Student’s t-test. Error bars=s.e.m. Conditions sharing letters are not statistically different from one another, one-**(F,G)** or two-way **(A,B)** ANOVA with Tukey’s post-hoc test. For all developmental rate experiments see Table S1 for full sample sizes and statistical results.

The microbiota-dependent growth delay of *Arc1* mutants motivated us to examine *Arc1* expression in the larval brain and gut under GNO and GF conditions. *Arc1* transcripts decreased slightly but significantly in the brains of GF compared to GNO larvae, but were unaffected in the larval gut (Fig. 2C). In the larval brain, there are multiple clusters of Arc1 positive neurons and other Arc1-positive cells in the central brain lobes (Mattaliano et al., 2007; Mosher et al., 2015; Fig. 2D,S3). In contrast to transcript levels, the number of Arc1-positive cells in the central brain lobes increased in GF *vs.* GNO animals (Fig. 2E,S3).

We next asked whether the polymicrobial GNO community’s ability to promote *Arc1* mutant growth rate was due to a single bacterial taxon or to the collective effects of the community. To test this, we mono-associated wild-type and *Arc1* null larvae with each of the four bacteria and assessed time to pupariation. Association with either of the two *Lactobacillus* isolates or *A. pasteurianus* accelerated both wild-type and *Arc1^E8^* larval growth, but was not sufficient to promote growth like the GNO community (Fig. 2F,G). In contrast, *Acetobacter sp.* was indistinguishable from GNO in promoting growth rate of both wild-type and *Arc1^E8^* animals (Fig. 2F,G). We did not find significant differences in larval bacterial loads among the four isolates that might explain the growth-promoting action of *Acetobacter sp.* in either genotype (Fig. S4A).

Because *Acetobacter sp.* alone was sufficient to achieve a growth rate comparable to wild-type in *Arc1^E8^* larvae, we asked whether it would also be necessary in the four-species GNO community. While mutant larvae associated with *A. pasteurianus*, *L. brevis*, and *L. plantarum* individually developed substantially slower than those associated with the four-species group or *Acetobacter sp.*, the three together in a GNO community lacking *Acetobacter sp.* were sufficient to promote normal development (GNO-3spp; Fig. 2G). For experimental tractability, we focused all subsequent investigation on larvae mono-associated with *Acetobacter sp.*.

While the delayed pupariation rate of wild-type GF larvae reflects a moderately extended duration of all three larval instars (Fig. S4B; Storelli et al., 2011), *Arc1^E8^* GF larvae undergo prolonged L1 and L2 phases (Fig. S4B). Further, GF animals of both genotypes increase in size at a more gradual rate than those with *Acetobacter sp.*, but this effect is exaggerated in *Arc1* mutants, suggesting a longer time to attain the critical weights that trigger larval molts and metamorphosis (Fig. S4C; Mirth et al., 2005). The extended larval period could reflect reduced nutrient intake, but we did not observe differences in feeding behavior or food consumption (Fig. S4D,E).

### Diet modulates the *Arc1*-dependent GF larval developmental delay

We have demonstrated that *Arc1* loss significantly exacerbates the developmental delay of GF larvae. The developmental rate of wild-type GF larvae is particularly sensitive to dietary yeast, the primary source of ingested amino acids, and also to food carbohydrate content (Shin et al., 2011; Storelli et al., 2011; Wong et al., 2014). To determine how *Arc1* deletion impacts microbe-dependent larval dietary sensitivity, we monitored time to pupariation of *Acetobacter sp.*-associated and GF wild-type and *Arc1^E8^* larvae fed nine yeast-dextrose diets of varying protein:carbohydrate ratios (Fig. 3; Lesperance and Broderick, 2020).

**Fig. 3.**
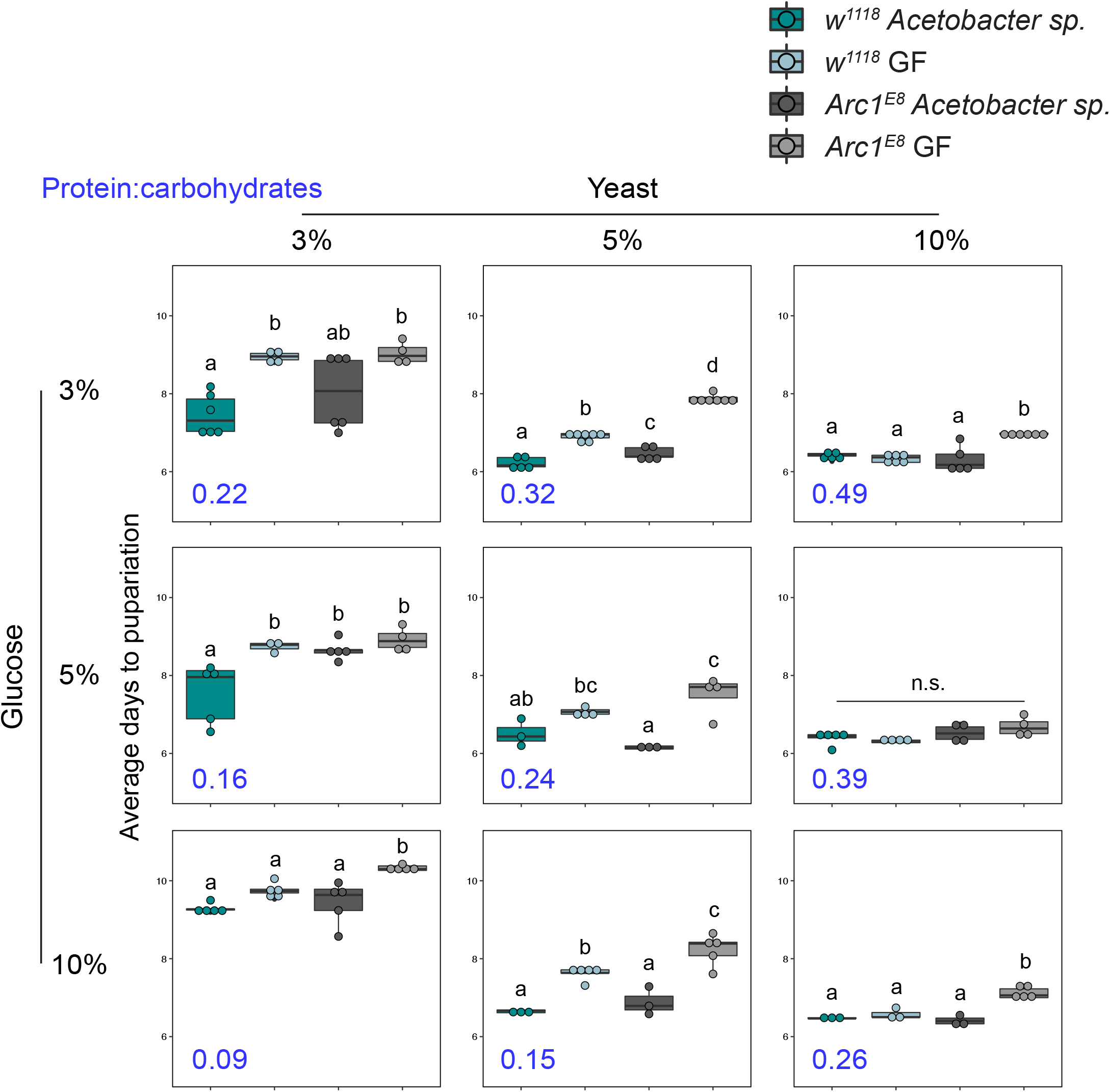
Host diet impacts microbial effects on wild-type and *Arc1* mutant developmental rate. Time to pupariation for wild-type and *Arc1^E8^ Acetobacter sp.*-associated and GF larvae reared on diets consisting of the indicated concentrations (weight/volume) of yeast and dextrose. Values in blue represent the protein:carbohydrate ratio for each diet as calculated with the Drosophila Diet Composition Calculator (Lesperance and Broderick, 2020a). Conditions that share a letter are not statistically different from one another, n.s.=not significant, two-way ANOVA with Tukey’s post-hoc test.

As expected, in wild-type animals “high-yeast” (10%) diets eliminated the developmental gap between *Acetobacter sp.*-associated and GF larvae, regardless of glucose concentration, while on “low-yeast” (3%) diets wild-type GF larvae were consistently delayed. The exception to this was the 3% yeast-10% glucose diet, which substantially slowed the developmental rate of all conditions, a known effect of high-sugar, low-protein diets (Musselman et al., 2011; Wong et al., 2014). Interestingly, on 5% yeast diets, increasing glucose concentration moderately accelerated the development of wild-type GF larvae, with a delay observed only at the lowest glucose level.

GF larvae lacking *Arc1* generally showed enhanced developmental sensitivity to dietary composition; GF *Arc1^E8^* animals developed more slowly than GF wild-type on 5/9 diets tested, including 2/3 “high yeast” diets (Fig. 3). On most diets, *Acetobacter sp.-* associated *Arc1* mutants developed at the same rate as *Acetobacter sp.*-associated wild-type larvae. One exception was the 3% yeast-5% glucose diet, where *Acetobacter sp.* failed to have any growth rate-promoting activity for *Arc1^E8^* larvae. Importantly, we did not find differences in *Acetobacter sp.* abundance on these diets that might correspond to the differences in growth rate of larvae of either genotype (Fig. S5).

While GF *Arc1*-deficient larvae exhibited developmental delay on multiple diets, none of those tested yielded a magnitude of delay comparable to that on our laboratory’s routine diet (P:C ratio 0.06; Fig. 2A). This may indicate that the full growth rate promoting effects of *Acetobacter sp.* require the more complex cornmeal-yeast-molasses diet and are less dependent on the protein:carbohydrate ratio.

Together, these results indicate that the importance of *Arc1* function during GF larval development is dependent on the host’s nutritional environment, and suggest that *Arc1* may be particularly important for GF larval growth dynamics in response to amino acid availability.

### Live *Acetobacter sp.* populations are required for optimal *Arc1* mutant growth rate

We next asked how *Acetobacter sp.* enables *Arc1*-deficient animals to develop at the same rate as wild-type mono-associated animals. We hypothesized that consumed bacteria may be a supplemental food source supporting *Arc1^E8^* development, as reported in wild-type *Drosophila* (Bing et al., 2018; Keebaugh et al., 2018; Storelli et al., 2017). To test this, we inoculated GF wild-type and *Arc1^E8^* cultures with heat-killed *Acetobacter sp.* cells daily throughout the larval growth period. This had no significant effect on GF wild-type or *Arc1* mutant larval developmental rates (Fig. 4A), providing no evidence for *Acetobacter sp.* cells as a nutritional supplement that affects growth.

**Fig. 4.**
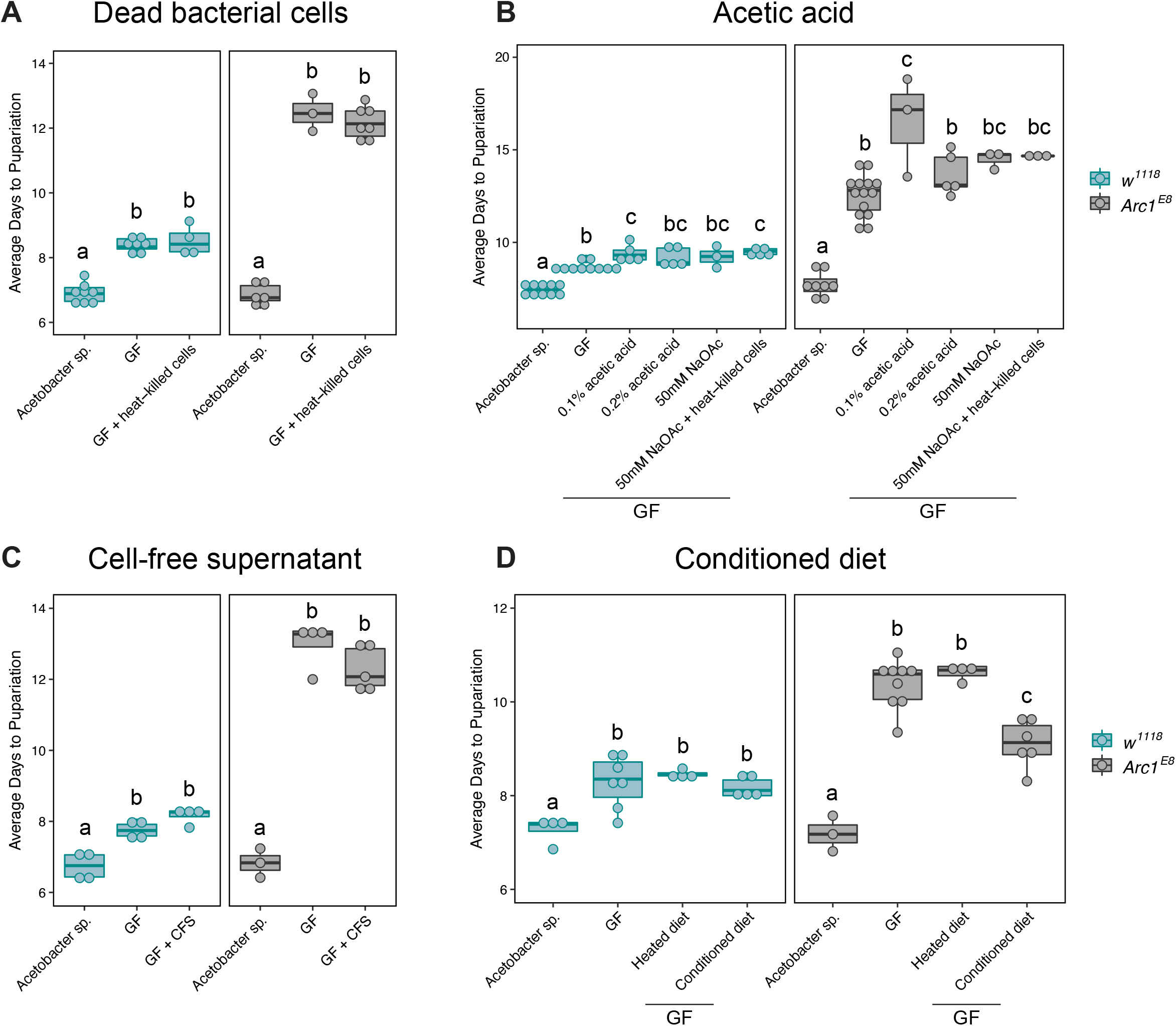
*Acetobacter sp.*-conditioned diet accelerates the development of *Arc1* mutants. **A.** Daily administration of heat-killed *Acetobacter sp.* planktonic culture has no effect on the developmental rate of GF wild-type or *Arc1^E8^* larvae. **B.** Rearing GF wild-type and *Arc1^E8^* larvae on diets containing acetic acid either further extends or has no effect on the larval developmental delay. **C.** Daily administration of filtered supernatant from *Acetobacter sp.* planktonic culture has no effect on developmental rate of GF wild-type or *Arc1^E8^* larvae. **D.** Rearing GF larvae on a sterile diet that has been pre-conditioned with *Acetobacter sp.* for five days (Conditioned diet) has no effect on wild-type, but partially restores the developmental rate of *Arc1^E8^* animals. Heated diet: GF diet heated under the same conditions used to kill *Acetobacter sp.* after pre-conditioning. Conditions that share a letter are not statistically different from one another, each panel analyzed by one-way ANOVA with Tukey’s post-hoc test.

The *Acetobacteraceae* generate acetic acid (Saichana et al., 2015), and two studies showed that acetic acid/acetate consumption can accelerate the development of GF larvae or larvae associated with an acetic acid-deficient *Acetobacter* (Kamareddine et al., 2018; Shin et al., 2011). Acetic acid has also been reported to yield no effect on growth, or to exacerbate the GF developmental delay at higher concentrations (Kim et al., 2017; Shin et al., 2011). Consistent with the latter, we found that providing GF wild-type or *Arc1^E8^* larvae with dietary acetic acid either had no impact on growth rate, or further slowed larval development (Fig. 4B). Feeding GF larvae acetic acid-supplemented diet in combination with heat-killed cells also had no effect (Fig. 4B). Additionally, we found that daily inoculation with filtered supernatant from planktonic *Acetobacter sp.* cultures had no effect on either wild-type or *Arc1^E8^* GF larvae (Fig. 4C), suggesting *Acetobacter sp.*-derived metabolite(s) do not promote growth.

*Drosophila* bacterial commensals proliferate on the flies’ diet and are continually ingested (Ludington and Ja, 2020). The host and bacteria therefore share a dietary niche, with the bacteria utilizing the flies’ food as a carbon source (Blum et al., 2013; Lesperance and Broderick, 2020b; Martino et al., 2018; Storelli et al., 2017). Certain microbe-dependent *Drosophila* traits, including growth rate, arise from bacterial utilization of dietary nutrients, producing an altered nutritional intake for the host (Consuegra et al., 2020; Huang and Douglas, 2015; Lesperance and Broderick, 2020b; Storelli et al., 2017). We therefore hypothesized that dietary modification might contribute to *Acetobacter sp.* support of *Arc1* mutant development. To test this, we inoculated GF food vials with *Acetobacter sp.*, allowed colonization of the diet for 5 days, and then killed all bacteria by heat treatment, resulting in an *Acetobacter sp.*-conditioned but microbiologically sterile food substrate.

The conditioned diet did not impact the developmental rate of wild-type GF larvae (Fig. 4D). In contrast, the conditioned diet substantially accelerated GF *Arc1^E8^* larval development, though these animals still developed significantly slower than *Arc1* mutants associated with live *Acetobacter sp.* (Fig. 4D). These data suggest that *Acetobacter sp.* modification of the larval diet is an important, though not exclusive, mechanism by which *Acetobacter sp.* promotes *Arc1^E8^* larval growth.

### *Acetobacter sp.* ameliorates systemic metabolism-related *Arc1* mutant phenotypes

We next asked whether the microbial environment impinges on other hallmarks of aberrant metabolism observed in *Arc1* mutants. Specifically, we assayed four organismal and cellular traits frequently associated with a larval growth delay and systemic metabolic dysregulation. In addition to slowed development, GF rearing stunts systemic growth yielding reduced pupal size (Fig. 5A; Kamareddine et al., 2018; Storelli et al., 2017). This size reduction was exaggerated in GF *Arc1* mutants (Fig. 5A). Microbiota perturbation can also compromise cellular growth, as measured by cell size in the adult wing (Shin et al., 2011). Trichome density in adult wings was increased in GF *Arc1^E8^* adults, indicating reduced cell size compared wild-type animals of either microbial condition (Fig. 5B). These data are consistent with impaired systemic and cellular growth capacity upon loss of both *Arc1* and the microbiota.

**Fig. 5.**
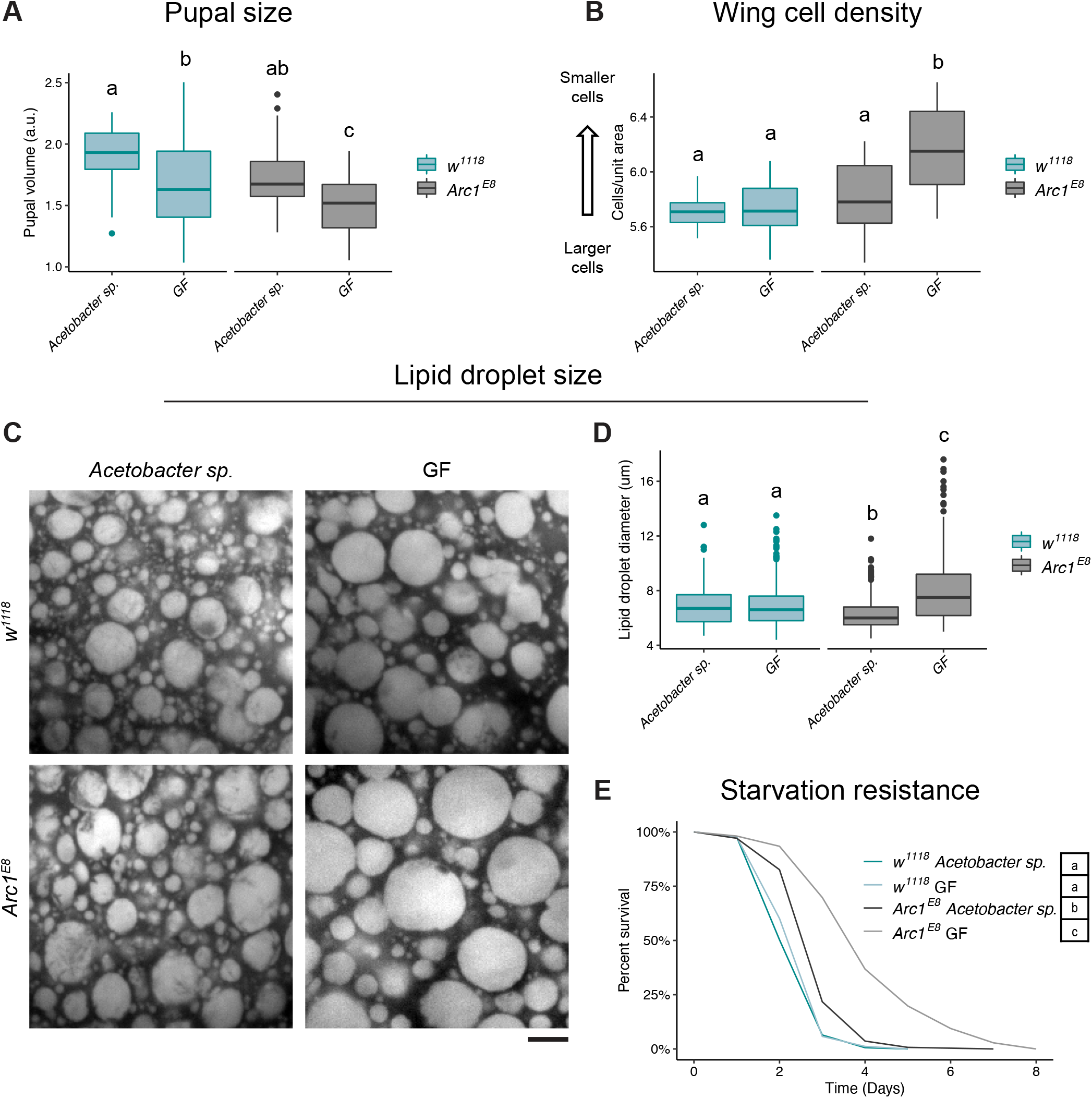
GF *Arc1* mutants exhibit additional growth and metabolic defects that are suppressed by mono-association with *Acetobacter sp.* **A.** Pupal volume is significantly decreased in GF *Arc1^E8^* animals. n=34-63 pupae per condition. **B.** Wing trichome density is increased (indicative of smaller cells) in GF *Arc1^E8^* adult females. n=13-20 wings per condition. **C,D.** Lipid droplet size is decreased in fat bodies of *Acetobacter sp.*-associated *Arc1^E8^* larvae and increased in GF *Arc1^E8^* larvae compared to wild-type animals under either microbial condition. n=407-512 lipid droplets from 15-20 animals per condition. Only droplets ≥5 µm were scored. Scale bar: 5 μm. E. *Arc1^E8^* mutants are more starvation resistant than wild-type, and this is significantly enhanced in GF *Arc1^E8^*. Conditions that share a letter are not statistically different from one another, Kruskal-Wallis test with Dunn’s multiple comparison (**A**,**B**,**D**), Cox proportional-hazards model analysis (**E**).

*Arc1* mutant larvae were previously shown to have elevated larval fat levels, and are starvation resistant as adults, the latter likely in part due to greater lipid stores accumulated during the larval period (Mattaliano et al., 2007; Mosher et al., 2015). Consistent with these reports, we found that GF *Arc1* mutants had significantly larger lipid droplets in the larval fat body (the primary adipose tissue; Fig. 5C,D), and adult females survived full nutrient deprivation for ∼2 days longer than wild-type adults (Fig. 5E).

Importantly, we primarily observed the effects of *Arc1* loss when *Arc1* mutants were reared GF; all four phenotypes were suppressed or altered when *Arc1* mutants were grown with *Acetobacter sp*. (Fig. 5A-E). Interestingly, *Acetobacter sp.*-associated *Arc1^E8^* larvae had significantly smaller lipid droplets in the fat body compared to wild-type animals (Fig. 5D). Comparable to published results (Mattaliano et al., 2007), *Acetobacter sp.*-associated *Arc1* mutants were more starvation resistant than wild-type, but succumbed considerably faster than GF mutants (Fig. 5E).

Collectively, these results indicate that loss of the microbiota uncovers or exacerbates growth and metabolic phenotypes in *Arc1*-deficient *Drosophila*, and association with a single bacterial taxon can strongly mitigate these defects.

### Selective *Arc1* expression in neuroendocrine cells mitigates growth and metabolic defects

We next investigated the tissues in which Arc1 function is important for regulating growth rate and other metabolism-related traits in GF animals. To this end, we selectively restored wild-type Arc1 in GF *Arc1^E8^* flies using a *UAS-Arc1* transgene (Mattaliano et al., 2007) and a variety of GAL4 drivers expressed in organs/cell-types with known roles in growth regulation and/or where Arc1 is endogenously expressed.

Weak ubiquitous (*armadillo-GAL4*) expression of *Arc1* significantly accelerated development in GF *Arc1^E8^* larvae, while expression in the anterior midgut, hindgut, and malpighian tubules (*drumstick-GAL4*), somatic muscle (*C57-GAL4*), and fat body (*r4-GAL4*) had no effect (Fig. 6A,S6A-D). Interestingly, pan-neuronal *Arc1* expression (*Appl-GAL4*) or strong expression in the foregut and midgut (*mex1-GAL4*) further extended the growth delay in GF mutants (Fig. 6A,S6E,F).

**Fig. 6.**
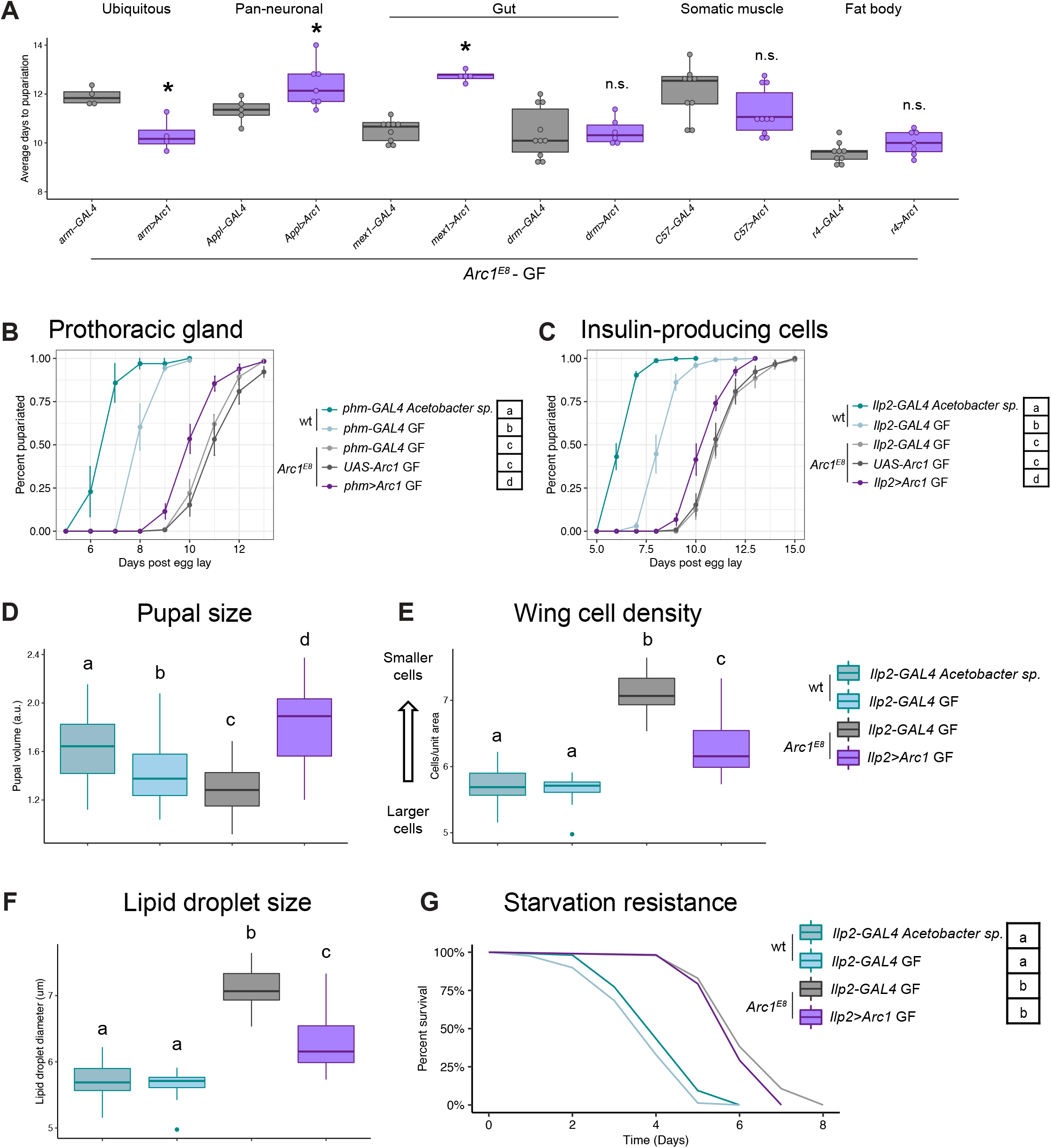
Arc1 expression in multiple tissues impacts growth and metabolic traits. **A.** Time to pupariation for GF *Arc1^E8^* larvae with *Arc1* expression selectively restored in the indicated tissues. Individual points represent biological replicates. *p<0.05, n.s.=not significant, see Figure S6 for full developmental rate growth curves and analyses. **B,C.** Selectively expressing *Arc1* in the prothoracic gland (**B**) or the insulin producing cells (IPCs, **C**) partially suppresses the *Arc1^E8^*-GF developmental delay. **D.** *Arc1* expression in the IPCs increases pupal size of *Arc1^E8^*-GF animals even beyond the wild-type controls. n=42-55 pupae per condition. **E.** *Arc1* IPC expression reduces increased trichome density (increases cell size) in *Arc1^E8^*-GF adult female wings. n=10-44 wings per condition. **F.** *Arc1* IPC expression reduces lipid droplet size in the *Arc1^E8^*-GF larval fat body. n=489-673 lipid droplets from 12-16 animals per condition. **G.** *Arc1* IPC expression does not affect starvation resistance under GF conditions. Conditions that share a letter are not statistically different from one another, one-way ANOVA with Tukey’s post-hoc test **(B,C,E)**, Kruskal-Wallis test with Dunn’s multiple comparison **(D,F)**, Cox proportional-hazards model analysis **(G)**.

Arc1 expression has been previously described in the ring gland (Mattaliano et al., 2007), a neuroendocrine organ which controls the timing of larval molts and pupariation through ecdysone biosynthesis (Mirth et al., 2005; Uryu et al., 2018; Yamanaka et al., 2015). Interestingly, Arc1 strongly increased in the ring gland once larvae stopped feeding and starting wandering in preparation for pupariation (Fig. S7A-B’’). Consistent with Arc1 playing a functional role in this growth-promoting tissue, selectively expressing Arc1 in the prothoracic gland (PG) cells of the ring gland moderately but significantly suppressed the growth delay of GF *Arc1* mutants (Fig. 6B).

Larval growth is also systemically regulated by the production of insulin-like peptides (Ilps) in the insulin-producing cells (IPCs) of the larval brain (Géminard et al., 2009; Rulifson et al., 2002). In wild-type animals, clusters of strongly Arc1-positive neurons are lateral to the medial IPCs (Fig. 2D,S7C-D). Weakly Ilp2-positive cells sometimes could be observed adjacent to those Arc1+ neurons (Fig. S7D, asterisk). Interestingly, in some cases, certain IPCs were weakly Arc1-positive (Fig. S7E-E’’, arrowheads), and we found weaker Arc1-positive cells adjacent to the IPCs (Fig. S7F-F’’, arrowheads). Overexpressing *Arc1* in the IPCs of GF *Arc1* null larvae (Fig. S7G-G’’) significantly increased larval growth rate, but, as with selective expression in the PG, not to wild-type levels (Fig. 6C). Expressing *Arc1* in the IPCs also suppressed the reduced pupal size (Fig. 6D), reduced wing cell size (Fig. 6E), and enlarged lipid droplets (Fig. 6F), but did not alter the starvation resistance of GF *Arc1* mutants (Fig. 6G). Notably, ubiquitous, IPC-, and PG-specific overexpression of Arc1 had no effect on developmental rate or growth in GF wild-type larvae (Fig. S8A-D), whereas pan-neuronal and foregut-midgut overexpression exacerbated the GF developmental delay, similar to our observations in the *Arc1^E8^* mutant background (Fig. S8E,F). Taken together, these data suggest that *Arc1* expression in multiple tissues is necessary to control larval growth and other metabolic traits.

### Evidence for microbe-dependent and -independent changes in insulin and ecdysone pathways in Arc1-deficient larvae

We next sought to identify molecular changes in *Arc1* null animals that might yield mechanistic insight into their growth and metabolic defects. Collectively, the phenotypes we observed in GF *Arc1^E8^* animals (Fig. 5) are reminiscent of those in IIS pathway mutants (Brogiolo et al., 2001; Broughton et al., 2005; DiAngelo and Birnbaum, 2009; Rulifson et al., 2002). For example, ablation of IPCs or loss of IIS signaling slows larval growth rate in GF or CV larvae (Fig. S9A; Rulifson et al., 2002). Surprisingly, we found that overactivation of IIS signaling in GF larvae through overexpression of Ilp2 or expression of activated PI3K also slows larval growth rate (Fig. S9B,C), similar to what was previously described for overexpression of Ilp8 (Vallejo et al., 2015). Previous work did not reveal altered IIS function in *Arc1* mutants (Mattaliano et al., 2007; Mosher et al., 2015), though under different dietary conditions and without microbial manipulation (see Discussion). Notably, *Acetobacter* strains sustain larval growth by activating IIS in wild-type flies fed nutrient restrictive diets (Kamareddine et al., 2018; Shin et al., 2011; Storelli et al., 2011). We therefore hypothesized that IIS function might be perturbed in a microbe-dependent manner in *Arc1^E8^* larvae.

In adequately fed larvae, insulin-like peptides (Ilps) are synthesized in the IPCs and secreted into the hemolymph; decreased *Ilp* expression and Ilp retention in the brain are associated with nutritional deficiency and impaired growth promotion (Géminard et al., 2009). We found that *Ilp3* and *Ilp5*, but not *Ilp2*, transcripts were reduced ∼2 fold in the brains of feeding GF *Arc1^E8^* larvae compared to wild-type larvae (Fig. 7A). By contrast, Ilp2 and Ilp5 protein levels were increased in *Acetobacter sp.*-associated *Arc1* mutant brains (Fig. 7B). Such accumulation is generally interpreted as evidence of decreased secretion and reduced IIS signaling. Interestingly, this retention was reduced in GF mutants (Fig. 7B).

**Fig. 7.**
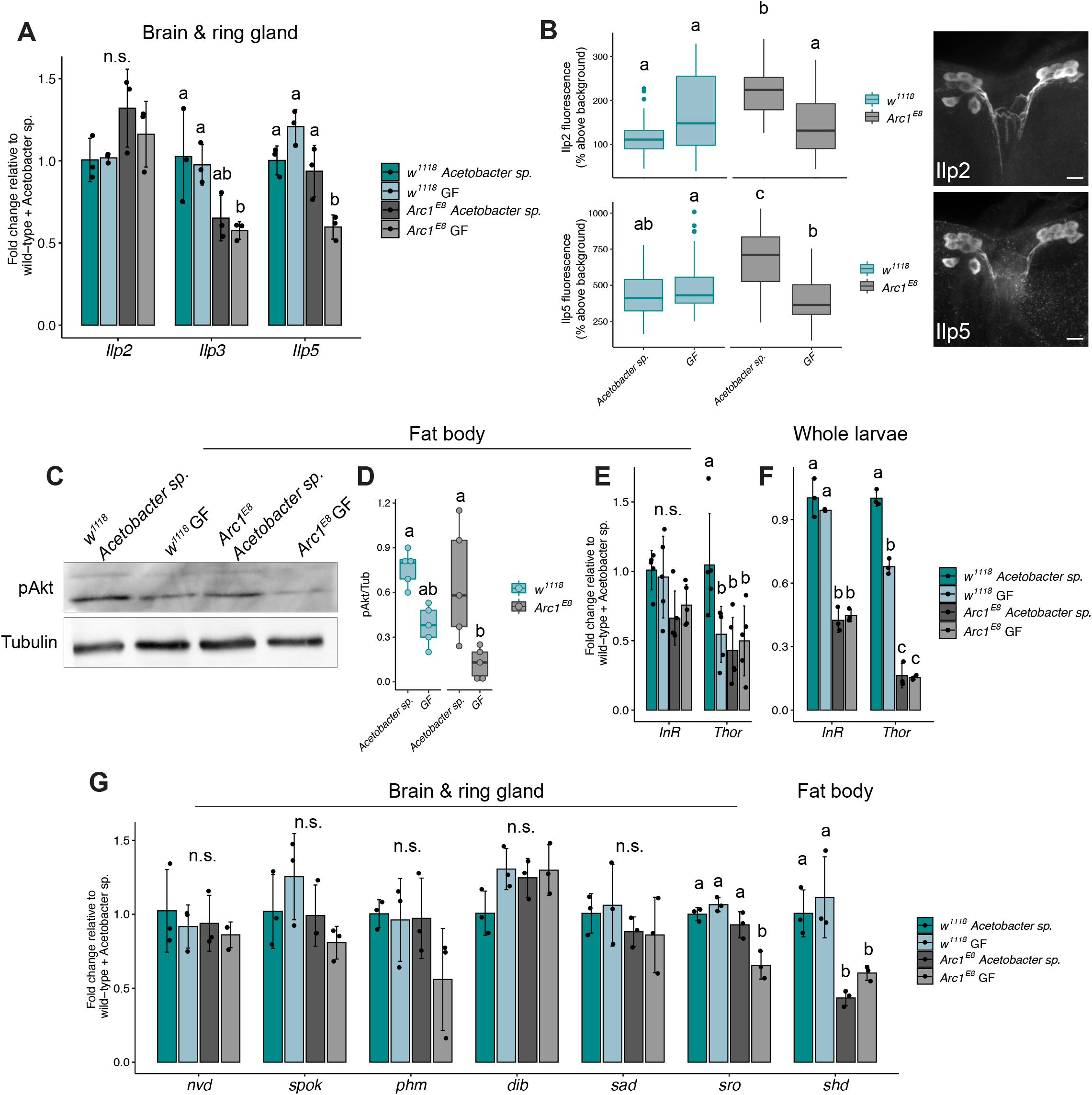
Evidence of disrupted insulin and ecdysone signaling in GF *Arc1* mutants. **A.** RT-qPCR analysis of *insulin-like peptide* (*Ilp*) gene expression in the brains of feeding *Acetobacter sp.*-associated and GF wild-type and *Arc1^E8^* larvae. **B.** Ilp2 and Ilp5 protein levels are elevated in the insulin-producing cells (IPCs) of *Acetobacter sp.*-associated *Arc1^E8^* larvae and this is suppressed in GF animals. Representative images of Ilp2 and Ilp5 immunostaining in larval brain IPCs. Scale bar: 5 μm. **C,D.** Phospho-Akt levels are reduced in the fat bodies of GF *Arc1^E8^* larvae compared to *Acetobacter sp.*-associated mutant and wild-type animals. **E,F.** RT-qPCR of the insulin pathway target genes *InR* and *Thor/4EBP* in the fat bodies of **(E)** or whole **(F)** feeding third instar larvae. **G.** RT-qPCR of ecdysone biosynthetic genes in the brain/ring gland, and the monooxygenase enzyme shade (*shd*) in fat bodies of feeding larvae as indicated. Conditions that share a letter are not statistically different from one another, n.s.=not significant, two-way ANOVA with Tukey’s post-hoc test.

Ilp secretion leads to systemic insulin signaling activation in key metabolic tissues, including the fat body (Brogiolo et al., 2001; DiAngelo and Birnbaum, 2009). Activation of IIS in these tissues leads to phosphorylation and activation of Akt (pAkt), nuclear exclusion of the transcription factor Foxo, and consequent upregulation of its negative targets *InR* and *Thor/4EBP* (Baker and Thummel, 2007). In GF *Arc1^E8^* fat body lysates, pAkt was decreased compared to *Acetobacter sp.*-associated fat bodies, and trended lower than GF wild-type, though the difference was not significant (Fig. 7C,D). Despite reduced pAkt levels (consistent with reduced IIS signaling), *InR* and *Thor* expression was either decreased (consistent with elevated IIS signaling) or unaffected in *Arc1* mutant fat bodies and whole animals, regardless of microbial condition (Fig. 7E,F). Interestingly, we observed the same trends of decreased pAkt, and decreased Foxo target gene expression in wild-type GF animals (Fig. 7C-F).

While we found that multiple IIS signaling steps are impacted by microbial condition and/or loss of *Arc1*, these changes do not suggest simple increased or decreased IIS pathway activity in GF *Arc1^E8^* larvae. This might suggest that the combined effects of *Arc1* and microbiota loss on IIS function are more complex than the effects of either manipulation alone, and thus could involve dysregulation of other pathways that impinge on IIS to affect metabolism and growth. The steroid hormone 20E regulates the timing of larval maturation, and is also nutrient responsive and intersects with IIS signaling (Boulan et al., 2013; Buhler et al., 2018; Yamanaka et al., 2015). Notably, the 20E biosynthetic pathway begins in the PG, where targeted *Arc1* expression suppresses the growth delay of GF *Arc1^E8^* larvae (Fig. 6B). *Shroud*, a short-chain dehydrogenase/reductase which functions in this pathway in the PG (Niwa et al., 2010), was moderately decreased (∼1.5 fold) in feeding GF *Arc1^E8^* larval brain/ring glands, while transcripts of other ecdysone biosynthetic enzymes were unaffected (Fig. 7G). The monooxygenase *shade* converts ecdysone to bioactive 20E in the fat body (Petryk et al., 2003), and reduced *shade* expression delays pupariation, stunts growth, reduces *Ilp3* expression, and induces Ilp2 retention (Buhler et al., 2018). Interestingly, fat body expression of *shade* decreased ∼2 fold in *Arc1* mutants under both microbial conditions (Fig. 7G).

Collectively, these data suggest that GF *Arc1* mutants experience complex dysregulation of two primary endocrine axes that regulate growth and metabolism in *Drosophila,* and some, but not all, of these defects are mitigated by *Acetobacter sp*.

## DISCUSSION

Many physiological traits in animals are shaped by highly complex, poorly understood interactions between a host’s genotype and its microbiota. Here we report an unexpected connection between the host gene *Arc1* and commensal bacteria that affects metabolism and growth in *Drosophila*. We initially identified *Arc1* from a screen for transcripts showing differential abundance in the heads of GF *vs.* CV/GNO adult flies. Notably, *Arc1* was recently designated a “core” microbiome-response gene in *Drosophila*, as it consistently appears in published RNA-seq studies focused on the gut or whole animals (Delbare et al., 2020). Our study further reveals that microbiota-dependent *Arc1* mRNA and protein changes vary in direction and magnitude between the gut and the head/brain, and among different populations of Arc1-positive cells in the brain (Fig. 1,2,S1,3). Arc1 expression increases substantially following neuronal activation, similar to mammalian Arc (Guan et al., 2005; Mattaliano et al., 2007; Montana and Littleton, 2006), and this response to activity can be brain region-specific (Mosher et al., 2015), comparable to the microbiota-dependent cell type-specific variance we observed. Genetic manipulations that reduce IIS activity also yield tissue-specific differential *Arc1* expression (Musselman et al., 2018; Tain et al., 2021), as does rearing flies on a high-fat diet (Rivera et al., 2019), or under starved conditions (Fig. 1B).

Together this complexity suggests that: 1) Arc1 expression is highly sensitive to microbial and nutritional cues, and 2) Arc1 might play mechanistically distinct roles in different tissues and cell types, necessitating precise spatiotemporal regulation of its transcript and protein levels. Consistent with the latter idea, targeted tissue-specific expression of Arc1 in *Arc1^E8^* GF animals had a range of effects depending on the driver (Fig. 6,S6). Notably, overexpressing Arc1 pan-neuronally and in the foregut-midgut was deleterious, significantly exacerbating the GF growth delay of both wild-type and *Arc1* mutant larvae (Fig. 6A,S6E,F,S8E,F). While IPC-specific and PG-specific expression both accelerated GF *Arc1^E8^* development, neither fully rescued the defect to wild-type GF growth rates (Fig. 6B,C). Similarly, IPC rescue only partially suppressed the reduced wing cell size and increased lipid droplet size of GF *Arc1* mutants, but resulted in pupae that were larger than wild-type animals, while having no impact on starvation resistance, in contrast to previous findings (Fig. 6D-G; Mattaliano et al., 2007). These results suggest that metabolic and growth homeostasis require precise modulation of Arc1 levels in multiple tissues. Also, Arc1 may function in mechanistically distinct ways in different tissues, with varying outcomes for the specific phenotypes we assayed. For example, Arc1 transcript is abundant in wing imaginal discs (FlyAtlas; Leader et al., 2018), raising the possibility that Arc1 might have a cell-autonomous function in this tissue which contributes to wing cell size (Fig. 5B). To date, both *Drosophila* and mammalian Arc proteins have been studied primarily in the brain and neurons. Our work suggests that Arc proteins may play important roles in other organs and cell types, and this should be examined in vertebrate systems as well.

Despite its consistently-reported microbe-dependent transcript changes, evidence for a functional interaction between Arc1 and the microbiota had not been previously investigated. We found that loss of *Arc1* in GF *Drosophila* resulted in deleterious phenotypes at the molecular, cellular, and organismal levels all indicative of metabolic dysregulation and compromised growth (Fig. 2,5,7). Some of these findings are similar to previously reported *Arc1* mutant phenotypes, which suggested that Arc1 regulates systemic metabolism. Specifically, Mosher et al. (2015) showed that *Arc1^esm18^* larvae have elevated whole-animal fat levels, which is consistent with our finding that GF *Arc1^E8^* larvae have enlarged lipid droplets in the fat body (Fig. 5C,D), and Mattaliano et al. (2007) reported that *Arc1^esm18^* adults are starvation resistant, which we also observed in *Arc1^E8^* adults (Fig. 5E). Importantly, as with developmental rate, pupal size, and wing cell size (Figs. 2A,5A-B), we only observed *Arc1* phenotypes (or, in the case of starvation resistance, observed the strongest effect) when *Arc1* mutants were also GF; *Arc1^E8^* animals mono-associated with *Acetobacter sp.* were more similar to, if not indistinguishable from, wild-type animals in most of our assays. It should also be noted that our study and those mentioned above utilized considerably different diet formulations, and, in fact, we found that the microbe-dependent *Arc1* larval growth rate was extremely sensitive to diet (Fig. 3). These results highlight the need to consider diet and microbial condition as key factors that can interact strongly with host genotype to greatly influence phenotypic outcomes, in *Drosophila* and other model systems.

While Arc1 was previously shown to impact systemic energy metabolism (Mosher et al., 2015), and our study has expanded the range of *Arc1* mutant phenotypes to include organismal and cellular growth defects, the exact molecular mechanisms through which Arc1 exerts these effects is unknown. Mammalian Arc promotes endocytosis of AMPA receptors via direct interactions with endocytic machinery (Chowdhury et al., 2006; DaSilva et al., 2016; Wall and Corrêa, 2018), and both *Drosophila* Arc1 and mammalian Arc encode polypeptides that can self-assemble into mRNA-containing capsid-like structures, which are released in extracellular vesicles and taken up by recipient cells (Ashley et al., 2018; Erlendsson et al., 2019; Pastuzyn et al., 2018). Both of these mechanisms appear to be critical for Arc’s role as a master regulator of synaptic plasticity. Their potential contributions towards its metabolic and growth-promoting functions have not been investigated, but it is possible that distinct molecular functions dependent on cellular and tissue contexts may contribute to Arc1’s role in metabolism and growth regulation.

The molecular and cellular processes impacted by the *Arc1*-microbiota-diet axis that might result in metabolic and growth phenotypes are also unclear. In *Drosophila*, these processes are largely coordinated by the intertwined activities of IIS and 20E signaling, which are both affected by the microbiota (Bing et al., 2018; Kamareddine et al., 2018; Shin et al., 2011; Storelli et al., 2011). Notably, we found that molecular readouts of IIS function at multiple points in the pathway, and in multiple tissues, were altered in *Arc1* mutants, though in both microbiota-dependent and -independent ways, and not consistent with simple IIS over- or under-activation (Fig. 7A-F). In addition to changes in the IIS pathway, two of the enzymes required for 20E biosynthesis are decreased in *Arc1* mutants (Fig. 7G). We think it is likely that the interplay between microbial condition and *Arc1* mutation has complex and multifarious impacts on many growth-regulating cellular processes, each of which may be a primary consequence of Arc1 loss, or secondary to the metabolic defects induced by Arc1 and microbiota removal. Interestingly, mammalian Arc has also been linked to insulin and metabolism. Arc expression can be strongly induced in cultured human neuroblasts by exogenous insulin (Kremerskothen et al., 2002), and mice fed a high-fat diet, which induces insulin resistance and diabetic-like phenotypes, have suppressed Arc expression in the cerebral cortex and hippocampus (Chen et al., 2020; Mateos et al., 2009). Our data suggest that links between Arc proteins and insulin signaling might be evolutionarily conserved, and that the influence of the microbiota may be an integral component of this relationship.

One of the major findings of this study is that mono-association with a single member of the bacterial microbiota, *Acetobacter sp.*, was sufficient to fully or partially suppress most of the phenotypes observed in GF *Arc1* mutants. We further found that pre-conditioning the larval diet with *Acetobacter sp.* was sufficient to accelerate GF *Arc1^E8^* development, suggesting that dietary modification is a key element of *Acetobacter sp.* growth-promoting activity (Fig. 4D). There is precedent for similar mechanisms in wild-type *Drosophila* associated with other microbiota members. For example, on nutritionally poor diets, *L. plantarum* depletes the levels of sugars and branched-chain amino acids, and increases the levels of glycolysis and fermentation products to promote larval growth (Storelli et al., 2017). Importantly, we found that live *Acetobacter sp.* populations are required for full growth rate promotion of *Arc1* mutants. In wild-type flies, commensal bacteria promote growth through interactions between bacterial cell wall components and gut cells, including induction of intestinal peptidase expression by D-alanylated teichoic acids (Consuegra et al., 2020; Matos et al., 2017), and innate immune signaling activation by DAP-type peptidoglycan (Davoodi et al., 2018; Kamareddine et al., 2018). Notably, these two modes of microbiota activity are reminiscent of microbial influence on mammalian physiology. Bacterial breakdown of macro-nutrients like complex polysaccharides in the human gut has been linked to health and disease states (Cockburn and Koropatkin, 2016). Other functions, such as immune cell maturation and maintenance of gut epithelial architecture, appear to require bacterial cell-derived antigens (Sekirov et al., 2010).

The ability of *Acetobacter sp.* to largely restore metabolic and developmental homeostasis to *Arc1* mutant *Drosophila* is a representative example of a bacterial symbiont mitigating the deleterious effects of genetic aberrations in the host, a prevalent feature of animal-microbe interactions (Douglas, 2019; Lynch and Hsiao, 2019; Ussar et al., 2016). The potential to harness these phenotypic buffering functions has motivated increasing efforts to develop microbial therapies for human diseases with a genetic basis (Baruch et al., 2021; Davar et al., 2021; Helmink et al., 2019). However, the success of these endeavors will require a more thorough understanding of the highly complex, modular interactions among host genotype, microbial metagenomes, diet, and other factors. This study reveals a tractable system to explore how a single host gene and the microbiota converge on conserved cellular pathways to regulate nutritional responsiveness, metabolic health, and developmental growth.

## MATERIALS AND METHODS

### *Drosophila* stocks and diets

The following fly stocks were used: *w^1118^*, Canton-S, and Oregon-R are long-term lab stocks originally from the Bloomington Drosophila Stock Center (BDSC), Top Banana (kind gift from Dr. Michael Dickinson), *yw* (BDSC #1495), *w^1118^; Arc1^E8^* (kind gift from Dr. Vivian Budnik and Dr. Travis Thomson), *Arc1^esm113^* (BDSC #37531), *Arc1^esm18^* (BDSC #37530), *yw;Arc2^EY21260^* (BDSC #22466), *UAS-Arc1-WT* (BDSC #37532), *arm-Gal4* (BDSC #1561), *Appl-Gal4* (BDSC #32040), *Ilp2^96A08^-GAL4* (BDSC #48030; Jenett et al., 2012), *phm-GAL4* (BDSC #80577), *drm-GAL4* (BDSC #7098), *mex1-GAL4* (kind gift from Dr. Claire Thomas), *C57-GAL4* (BDSC #32556), *r4-GAL4* (BDSC #33832), *Hs-GAL4* (BDSC #1799), *UAS-Ilp2* (BDSC #80936), *UAS-rpr* (BDSC # 5824), *UAS-PI3K92E^CAAX^* (BDSC #8294). We use a yeast-cornmeal-molasses diet consisting of (percentages given as wt/vol or vol/vol throughout): 8.5% molasses (Domino Foods), 7% cornmeal (Prairie Mills Products), 1.1% active dry yeast (Genesee Scientific), 0.86% gelidium agar (MoorAgar), 0.27% propionic acid (Sigma), and 0.27% methylparaben (Sigma). All experiments utilized this diet except those in Fig. 3, which were conducted on yeast-glucose diets. Yeast-glucose diets were prepared as described (Koyle et al., 2016) and consisted of the indicated proportions of active dry yeast (Genesee Scientific) and dextrose (Fisher Scientific), with gelidium agar, propionic acid, and methylparaben as above.

### Bacterial stocks

The *Acetobacter sp.*, *Acetobacter pasteurianus*, *Lactobacillus plantarum*, and *Lactobacillus brevis* stocks utilized in this study were all isolated from conventionally-reared Top Banana *Drosophila* cultures in our laboratory. Adult flies were surface sterilized in 10% sodium hypochlorite and 70% ethanol, rinsed three times and homogenized in phosphate buffered saline (PBS). Serial dilutions of fly homogenates were plated on de Man, Rogosa, and Sharpe (MRS; Weber Scientific) and acetic acid-ethanol (AE; 0.8% yeast extract, 1.5% peptone, 1% dextrose, 0.5% ethanol, 0.3% acetic acid; Blum et al., 2013) agar plates. Colonies with distinct morphology were streaked for isolation. Bacterial taxonomies were assigned by PCR amplification and sequencing of the 16S rRNA gene using universal bacterial primers 8F (5’-AGAGTTTGATCTGGCTCAG-3’) and 1492R (5’-GGMTACCTTGTTACGACTT-3’; Eden et al., 1991). Sequences were searched against the NCBI nr/nt database via blastn (Altschul et al., 1990; Camacho et al., 2009; Morgulis et al., 2008) and taxonomies were assigned based on >97% sequence homology. Because the 16S rRNA sequence for isolate A22 (*Acetobacter sp.*) bore >97% similarity with >five different *Acetobacter* species we did not assign a species-level taxonomic classification.

### Generation of germ free and gnotobiotic fly cultures

Germ free and gnotobiotic *Drosophila* cultures were generated according to established methods (Koyle et al., 2016). Synchronous populations of embryos were collected on apple juice agar plates. In a sterile biosafety cabinet, embryos were treated with 50% sodium hypochlorite solution for 2 min to eliminate exogenous microbes and remove the chorion. Embryos were then rinsed twice in 70% ethanol, twice in sterilized milliQ water, and once in sterilized embryo wash (2% Triton X-100, 7% NaCl). Sterilized embryos were then pipetted into autoclaved food vials to generate germ free cultures. To generate gnotobiotic flies, overnight cultures of bacterial isolates were grown in MRS broth (30°C with shaking for *Acetobacter* isolates and 37°C static for *Lactobacilli*). Sterile food vials were inoculated with 40 μL of overnight cultures (OD∼1) immediately prior to the addition of sterilized fly embryos. For poly-associated (GNO) flies, vials were inoculated with 40 μL of a 1:1:1:1 mixture of overnight cultures of the four indicated bacteria.

For mono-association experiments (Fig. 2), bacterial inoculum was standardized to ∼10^8^ CFU for each strain tested using the following empirically determined CFU/mL constants: *Acetobacter sp.*: 4.5×10^8^, *A. pasteurianus*: 6.12×10^8^, *L. brevis*: 4.28×10^8^, *L. plantarum*: 1.14×10^8^ (Koyle et al., 2016).

All experimental *Drosophila* cultures were maintained in an insect incubator at 23°C, 70% humidity, on a 12:12 light-dark cycle.

Larval and adult animals were confirmed as germ free or mono-/poly-associated by homogenization in sterile PBS and plating on MRS and AE agar plates.

We did not maintain GF, GNO, or mono-associated flies over multiple generations; all experiments utilized independently-derived germ free or gnotobiotic animals.

### Developmental timing measurements

Synchronous populations of embryos were collected in a 4-6 hr time window and treated as described above to generate cultures of defined microbial conditions. For pupariation and eclosion rate analysis, the number of pupae formed or empty pupal cases, respectively, were counted daily until 100% of the population had pupariated or eclosed. The duration of larval development is strongly affected by crowding conditions in the food (Klepsatel et al., 2018). Also, variable and unpredictable numbers of embryos do not survive the bleach and ethanol washes employed to generate germ free and gnotobiotic cultures (Koyle et al., 2016; Troha and Buchon, 2019; unpublished observations). We found that 10-30 larvae per vial was optimal for growth, and that within-treatment developmental rates were comparable for vials with larval densities in this range (unpublished observations). Therefore, vials containing fewer than 10 and greater than 30 animals were omitted from analyses as either under- or over-crowded, respectively. Larval instars were determined via mouth hook and/or posterior spiracle morphologies (Oldroyd, 1951).

Larval length and pupal volume were measured from images using Fiji (Schindelin et al., 2012). For pupal volume, length (l) and width (w) of each pupa were measured and volume calculated as previously described (Layalle et al., 2008; Redhai et al., 2020): V=4/3*π*(l/2)(w/2)^2^.

### Larval feeding assays

Larval feeding was assessed via dye consumption (Buhler et al., 2018; Libert et al., 2007; Mosher et al., 2015; Shin et al., 2011). Pre-wandering third instar larvae were transferred to apple juice plates coated with a layer of yeast paste mixed with FD&C Red #40 dye (Ward’s Science) and allowed to feed for 30 min at 23°C. Guts were then dissected from 20 larvae, homogenized in 1 mL PBS, and absorbance of the supernatant was measured at 490nm with a Tecan Spark microplate reader.

Mouth hook contraction rates were assayed as described (Bhatt and Neckameyer, 2013). Pre-wandering third instar larvae were transferred to apple juice plates thinly coated with yeast paste, given 30 s to acclimate, and contractions were counted manually for 30 s.

### Quantifying bacterial loads for mono-associated larvae

Pre-wandering third instar larvae (8-10 animals per replicate) were removed from the food and surface sterilized in 10% sodium hypochlorite for 1 min. Larvae were then rinsed three times in PBS and homogenized in 125 μL PBS via bead beating for 30 s. Homogenates were serially diluted in PBS and dilutions were plated on AE (for *Acetobacter*) or MRS (for *Lactobacilli*) agar plates. Plates were incubated for 2-3 days at either 30°C (AE) or 37°C (MRS), and resultant colonies were counted manually from dilution plates bearing ∼50-400 colonies. Bacterial loads were calculated as colony-forming units (CFUs) per larva, as previously described (Koyle et al., 2016), and log transformed for analysis.

### Dietary treatments

#### Heat-killed bacterial feeding (Fig. 4A)

Overnight cultures of *Acetobacter sp.* were heat-killed at 65°C for 3 hr and autoclaved food vials were inoculated with 40 μL of heat-killed suspension prior to the addition of GF embryos. Vials were further inoculated with 40 μL heat-killed bacterial suspension daily until 100% of the population had pupariated. Successful heat killing was confirmed for each daily inoculum by plating undiluted heat-treated bacterial suspension on AE plates, and by plating larval homogenates.

#### Acetic acid-supplemented diets (Fig. 4B)

Fly food was autoclaved and allowed to cool to ∼50-60°C. Glacial acetic acid (Fisher Scientific) was added to 0.1% or 0.2%, or sodium acetate (Sigma) was added to a final concentration of 50 mM.

#### Cell-free supernatant feeding (Fig. 4C)

Overnight cultures of *Acetobacter sp.* (3 mL in MRS, grown as described above) were filtered twice through 0.22 μm PVDF sterile membrane filters (Genesee Scientific). Autoclaved food vials were inoculated with 40 μL of filtered media immediately prior to addition of GF embryos, and vials were further inoculated with 40 μL of filtered media daily until 100% of the population pupariated. Absence of live bacterial cells was confirmed by plating daily filtered media on AE plates, and plating larval homogenates.

#### *Acetobacter sp.*-conditioned diet (Fig. 4D)

Autoclaved food vials were inoculated with *Acetobacter sp.* overnight culture, as described above. Inoculated vials were incubated for five days at 23°C. Vials were then incubated at 65°C for 1 hr to kill bacteria. Sterility of the conditioned diet was confirmed by plating food and larval homogenates on AE plates. Heated diet controls consisted of un-inoculated, autoclaved GF vials incubated at 65°C for 1 hr.

### Wing analysis

Wings from adult females were dissected, mounted in Aqua-mount (Thermo Scientific), and imaged with a QICAM-IR Fast 1394 camera (Q-Imaging) on a Zeiss Axioskop2 Plus microscope. Trichome density was measured with FijiWings2.3 software using the 150px trichome density feature (Dobens and Dobens, 2013).

### Starvation resistance

Starvation experiments were conducted by transferring 8-10 five-day old adult females to 1% agar-water vials. ∼50-120 animals per condition per replicate were assessed. Flies were transferred to fresh agar-water vials daily and survival monitored daily until 100% of the population succumbed. Data from multiple replicates (2-3 per condition) were combined for analysis.

### RT-qPCR

Tissues were dissected in cold PBS and homogenized immediately in Trizol reagent (Thermo Fisher). RNA was extracted using the Direct-zol RNA Miniprep kit (Zymo Research) exactly following the manufacturer’s protocol. High quality RNA (A_260nm/280nm_ ∼2; 250-500 ng) was used as template for cDNA synthesis using the qScript cDNA synthesis kit (QuantaBio). Product from cDNA synthesis reactions was used for qPCR with the PerfeCTa SYBR Green Supermix (QuantaBio) in an Applied Biosystems 7300 Real Time PCR System instrument. Data were normalized to *Rpl32* and expression fold changes were calculated using the 2^-ΔΔCt^ method. Primers sequences are listed in Table S2.

### Immunohistochemistry

For immunostaining of adult and larval brains and adult guts, tissues were dissected in cold PBS and fixed in 4% paraformaldehyde-PBS (Electron Microscopy Sciences) for 30 min at room temperature. Samples were washed in PBS with 0.3% Triton X-100 (PBT) and blocked for 1 hr in PBS with 0.3% Triton X-100 and 5% normal goat serum (PNT). Incubations with primary antibodies in PNT were 4°C overnight with agitation. After washing in PBT, samples were incubated with secondary antibodies in PNT for 3 hr at room temperature. Samples were washed in PBS and mounted in aqua-poly/mount (Polysciences) or ProLong Gold antifade mounting medium (Invitrogen).

Primary antibodies: rabbit anti-Arc1 (1:250; Ashley et al., 2018), rat anti-Ilp2 and rabbit anti-Ilp5 (1:800; Géminard et al., 2009), mouse anti-EcR common (1:100; Developmental Studies Hybridoma Bank (DSHB) DDA2.7). Secondary antibodies: AlexaFluor488, AlexaFluor546, or AlexaFluor647 conjugated (Invitrogen) at 1:1000.

For lipid droplet analysis, larval fat bodies were dissected in PBS, transferred to 6 mm-well Shandon multi-spot slides (Thermo Scientific), and fixed for 30 min in 4% paraformaldehyde-PBS. Fat bodies were then rinsed three times in PBS, incubated with BODIPY 493/503 (1 mg/ml; Thermo Fisher) diluted 1:1000 in PBS for 30 min at room temperature, rinsed three times in PBS, and mounted as above.

### Image acquisition and analysis

Images were captured on a spinning disc microscope with a Celesta 1W light engine (Lumencor), an X-Light V2 scan head (Crest Optics), and a Prime95B CMOS camera (Photometrics) on a Zeiss Axiovert 200M using Metamorph software (Molecular Devices). All analyses were conducted using Fiji. Fluorescence intensity was calculated as the percent fluorescence above background. Lipid droplets *≥*5 µm were measured in a single z-plane that represented the largest diameter droplets between the tissue surface and the nuclei. Low magnification images of adult guts (Fig. 1E) were stitched together manually using Fiji.

### Western blots

Fat bodies from ten, pre-wandering third instar larvae were dissected in PBS, homogenized via bead beating in 180 μL ice cold PBS, and immediately frozen until analysis. Lysates were centrifuged to pellet tissue debris and equal volumes mixed 1:1 with 2x Laemmli sample buffer (BioRad). Samples were boiled, resolved on 10% SDS-PAGE gel, and transferred to 0.22 μm pore-size nitrocellulose membrane (BioRad). Blots were blocked in 5% bovine serum albumin in tris-buffered saline with 0.1% Tween 20 (TBS-T) and incubated overnight at 4°C with rabbit anti-phospo-Ser505-Akt (Cell Signaling Technology 4054) or mouse anti-*α*-Tubulin (DSHB 12G10) diluted 1:1000 in blocking buffer. Incubations with horseradish peroxidase-conjugated secondary antibodies (1:20,000; Jackson ImmunoResearch) were performed for 3 hr at room temperature, and signal detected with Pierce ECL substrate (Thermo Fisher) on a ChemiDoc MP imaging system (BioRad). Quantification of relative protein amounts was conducted using Fiji.

### Statistical analysis

Statistical tests were conducted and Figures generated using Prism 9 (GraphPad) and R version 3.5.1 (R Core Team, 2019). RT-qPCR and immunostaining data were analyzed via Student’s t-test for comparison of two groups, or via two-way ANOVA for comparison of genotype and microbial condition. For developmental rate data, the average time to pupariation was calculated for each vial from the number of individuals pupariating on each day until the entire population completed larval development. These per-vial values from at minimum three replicates were used for statistical analyses; full sample sizes and statistical test output for all development experiments are reported in Table S1. Within-genotype comparisons among different treatments were conducted via one-way analysis-of-variance (ANOVA), while comparisons among different genotypes and treatments were conducted via two-way ANOVA. Post-hoc analysis among significantly different factors were conducted via Tukey test. Data that did not meet parametric test assumptions (normal distribution assessed via Shapiro-Wilk test and homogeneity of variance assessed by Levene’s test) were analyzed via Mann-Whitney test or Kruskal-Wallis test with Dunn’s post-hoc comparison (Bonferroni correction). Tests used for each experiment are indicated in the Figure legends. Starvation survival data were compared via Cox proportional-hazards model analysis using the “survival” package (Therneau, 2012). Throughout, the threshold of statistical significance was considered p<0.05.

## Supporting information

Keith et al. Supplemental Figures

## ACKNOWLEDGMENTS

We thank members of the McCartney, Hiller, and Mitchell labs for helpful discussions while conducting the study. We thank Rory Eutsey (Hiller lab, Carnegie Mellon University) for technical assistance with qPCR experiments. We would like to thank Dr. John Woolford for providing feedback on the manuscript. The Top Banana fly stock was a generous gift from Dr. Michael Dickinson’s lab (CalTech). The *Arc1^E8^* fly stock and Arc1 antibody were generous gifts from Dr. Vivian Budnik and Dr. Travis Thomson (University of Massachusetts Medical School). *Mex1-GAL4* flies were a kind gift from Dr. Claire Thomas (Pennsylvania State University). The Ilp2 and Ilp5 antibodies were generous gifts from Dr. Pierre Léopold (Institut Curie). We thank the Bloomington Drosophila Stock Center for providing other fly stocks. We would like to thank the Woolford, Mitchell, and Hinman labs and the Molecular Biosensor and Imaging Center at Carnegie Mellon University for reagents and equipment.

