## Supplementary material for "Arc1 and the microbiota together modulate growth and metabolic traits in *Drosophila*": Keith et al. Supplemental Figures

### Supplementary figures

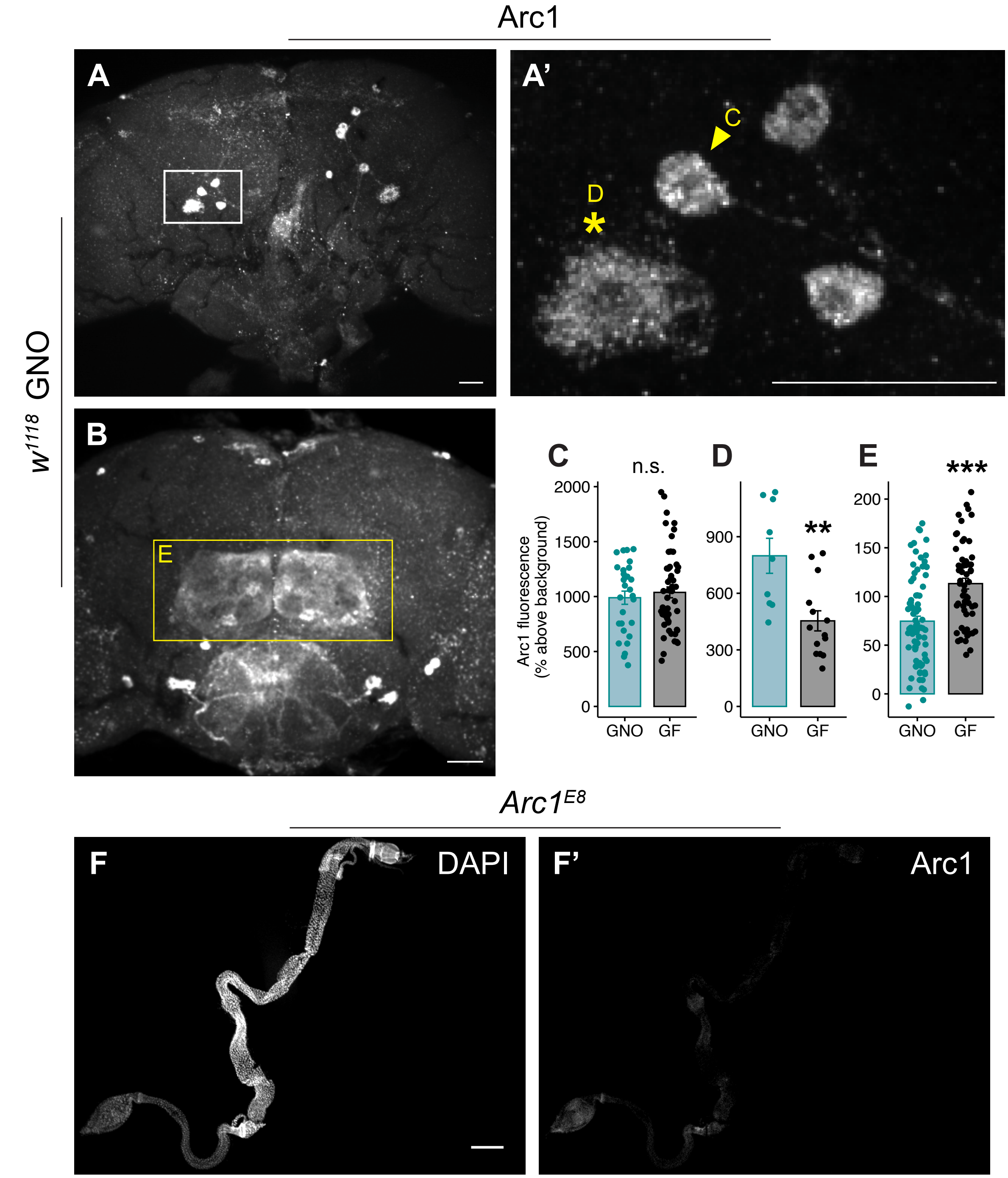


**Fig. S1. Arc1 immunofluorescence in adult tissues. A,A’.** Populations of Arc1-positive central brain cells with distinct Arc1 staining patterns in the posterior adult brain. **A’.** Higher magnification image of boxed region in **A**, showing clusters of central brain neurons with concentrated bright Arc1 staining (example cell indicated with arrowhead), and nearby cells with weaker, more diffuse Arc1 staining (indicated with asterisk). **B.** Arc1 expression pattern on the anterior surface of adult brains. Boxed area indicates the antennal lobes. Scale bars: 5 μm (**A,A',B**). **C-E.** Arc1 fluorescence intensity comparisons between GNO and GF animals for different brain cell types. **C.** Arc1 intensity in brightly-labeled central neurons of the posterior brain (indicated with arrowhead in **A'**) is the same in GNO and GF flies. **D.** In GF animals, there is reduced signal in cells showing more diffuse Arc1 staining in the posterior adult brain (indicated with asterisk in **A'**). **E.** Increased Arc1 fluorescence intensity in the antennal lobes (boxed region in **B**) in GF *vs.* GNO brains. All analyses were conducted with 5-6 day old adult male $w^{1118}$ flies. Individual points represent individual cells analyzed, **C,D** n=6-9 animals per condition. **E** n=7-8 animals per condition. **p<0.01, ***p<0.0001 n.s.=not significant, Mann-Whitney test (**C**) or Student's t-test (**D,E**). Error bars = s.e.m. **F,F'.** Arc1 staining in adult gut of *Arc1* null flies, showing minimal non-specific background staining. Scale bar: 200 μm.

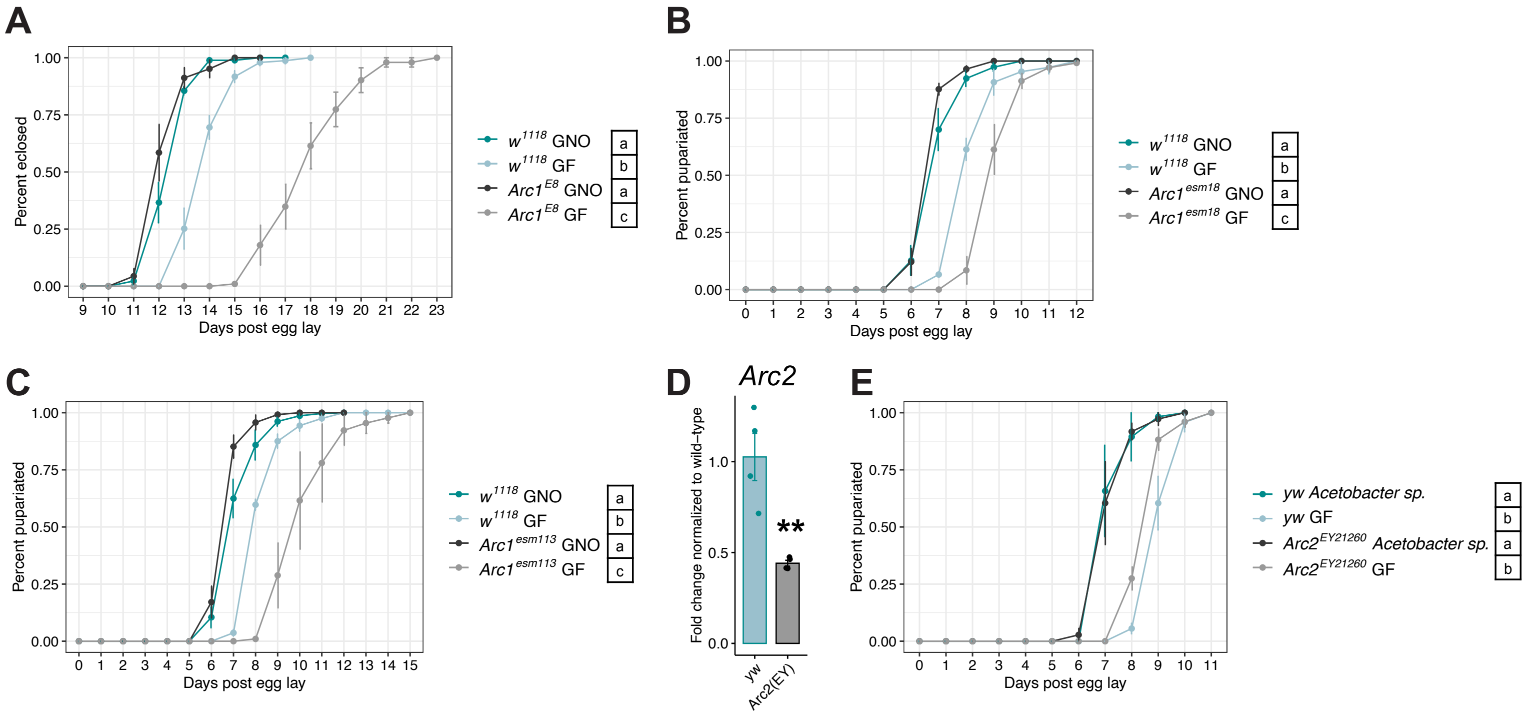


**Fig. S2. Additional *Arc1* mutants are developmentally delayed under GF but not GNO conditions. A.** Time to eclosion of wild-type and *Arc1* mutant populations represented in Fig. 2A. **B,C.** Time to pupariation for wild-type larvae and animals bearing independent *Arc1* mutant alleles. **D.** RT-qPCR of *Arc2* transcripts in whole pre-wandering third instar larvae of the indicated genotypes. $Arc2^{EY}$ is a random P-element insertion of $w^{+mC}y^{+mDint2}UASp$ in the 3' UTR of *Arc2*. Individual points represent normalized values for each biological replicate, 10 larvae per replicate. ** $p<0.01$, Student's t-test. **E.** Time to pupariation for wild-type and $Arc2^{EY}$ animals grown GF or in mono-association with *Acetobacter sp.* Reduction of *Arc2* does not impact time to pupariation. Error bars=s.e.m. For all developmental rate data **(A-C, E)**, conditions sharing a letter are not statistically different from one another, two-way ANOVA with Tukey's post-hoc test.

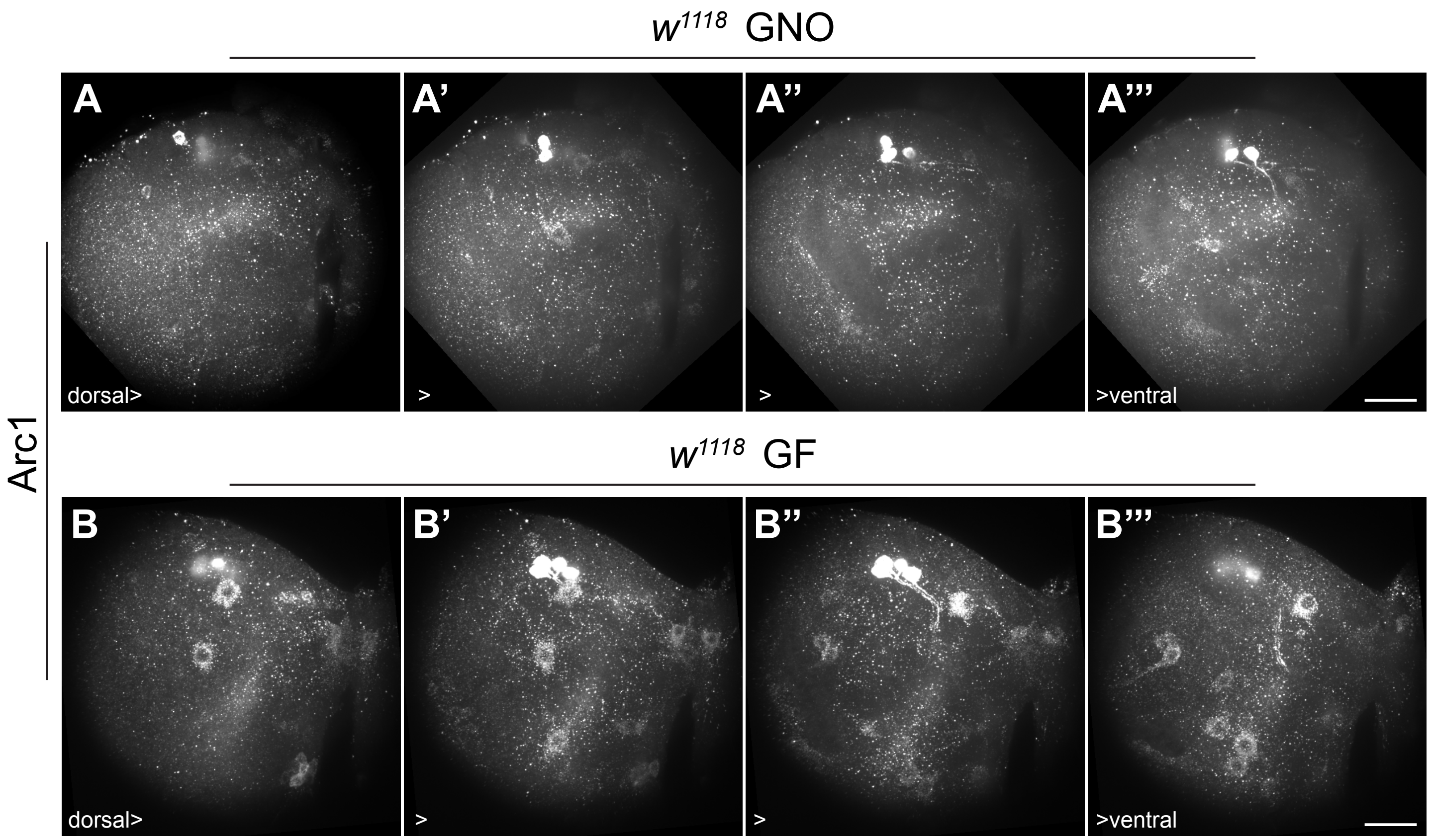


**Fig. S3. Increased number of Arc1-positive cells in brains of GF larvae compared to GNO larvae.** Arc1 staining in a representative lobe from each brain shown in Fig. 2D. Each image is a projection of eleven 0.2 µm slices. These are serial sections where the dorsal most sections are on the left and the ventral most sections are on the right. The signal is saturated in the highly expressing central brain neurons to more easily detect the other Arc1-positive cells that express at lower levels. Scale bar: 5 µm.

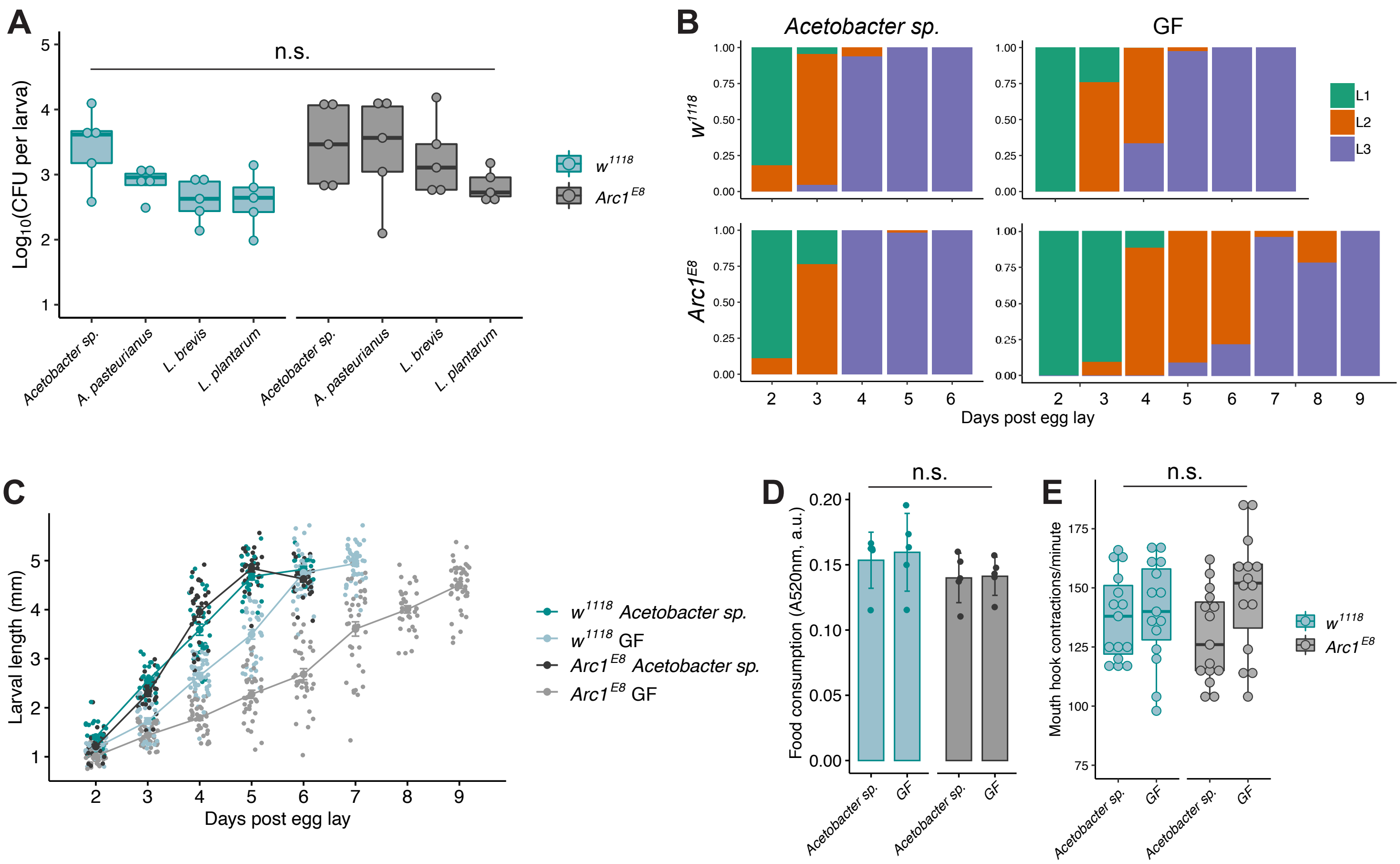


**Fig. S4. Characterization of microbial loads, larval growth, and feeding behavior. A.** Microbial loads for wild-type and *Arc1* mutant larvae mono-associated with the four bacterial isolates. Each data point represents a biological replicate of log-transformed colony forming unit (CFU) counts per larva recovered from surface-sterilized pre-wandering third instar larvae, 8-10 larvae per replicate. n.s.=not significant, two-way ANOVA with Tukey's post-hoc test. **B.** Percentage of larvae in the indicated instar stage daily until 100% of the population reaches the third instar stage. Data represent pooled percentages from three biological replicates for each day, ~30-60 animals per day. **C.** Larval length for both genotypes and microbial conditions over time. Each data point represents an individual larva, three biological replicates, ~10-20 larvae per replicate. **D.** Food consumed by larvae of each genotype and microbial condition in 30 min. Each data point represents food content in pooled guts of 20 pre-wandering third instar larvae. **E.** Mouth hook contraction rates, each point represents an individual larva. Error bars=s.e.m. n.s.=not significant, two-way ANOVA.

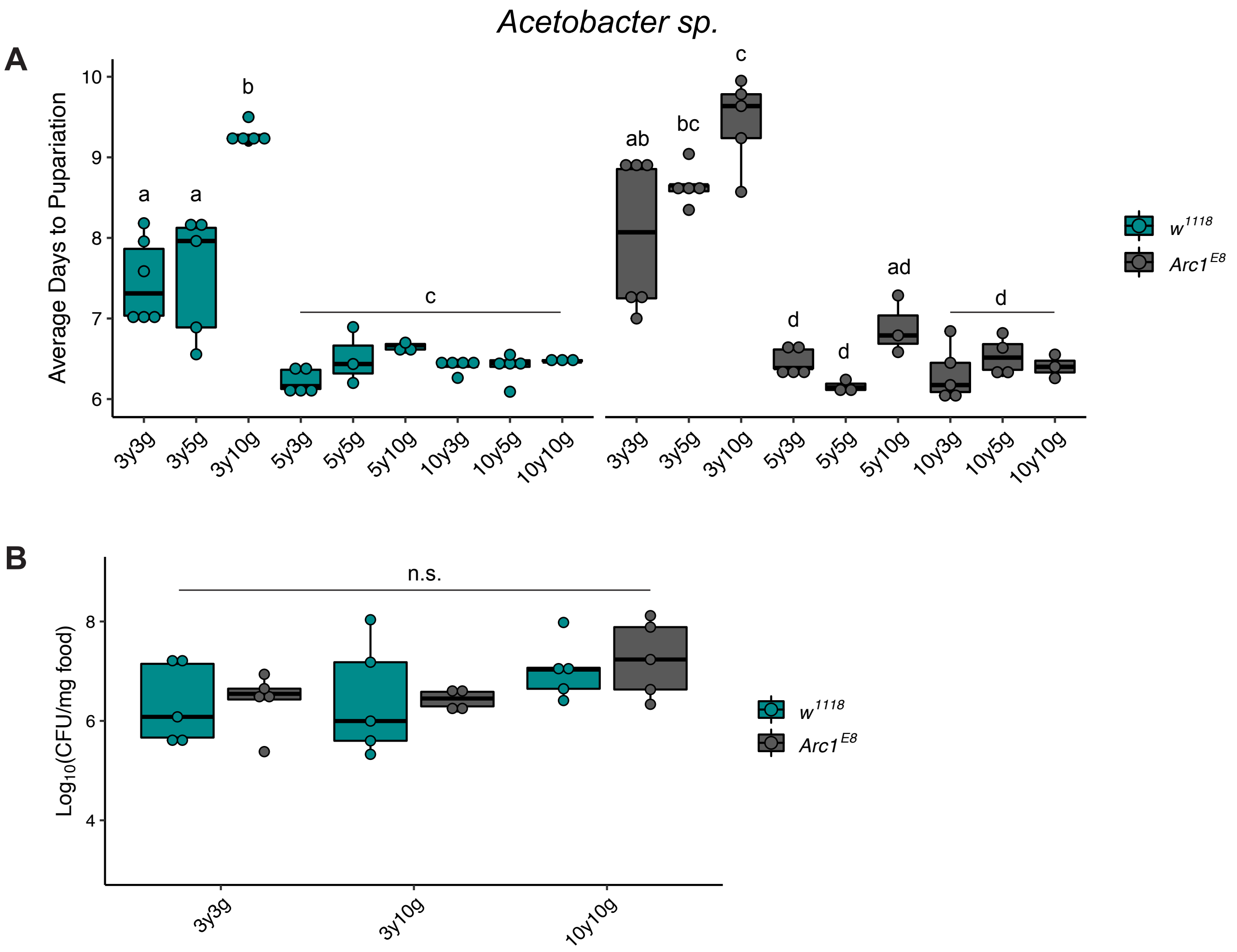


**Fig. S5. *Acetobacter sp.* colonization of yeast-glucose diets.** Testing the hypothesis that differences in *Acetobacter sp.*-associated larval growth rates among the various yeast-glucose diets **(A)** might result from varied *Acetobacter sp.*-colonization levels on these diets **(B)**. **A.** Developmental rate data for *Acetobacter sp.*-associated wild-type and *Arc1* mutant larvae grown on nine yeast-glucose diets. These are the same data presented in Fig. 3, but plotted collectively here for direct comparison of diet-dependent impacts on growth rate under *Acetobacter sp.*-associated conditions. **B.** *Acetobacter sp.* colonization levels on indicated diets. These three diets result in different growth rates for both genotypes, as shown in **A**., but *Acetobacter sp.* does not have significantly different levels of colonization on these diets. Conditions sharing a letter are not statistically different from one another, n.s.=not significant, each genotype compared by one-way ANOVA with Tukey's post-hoc test.

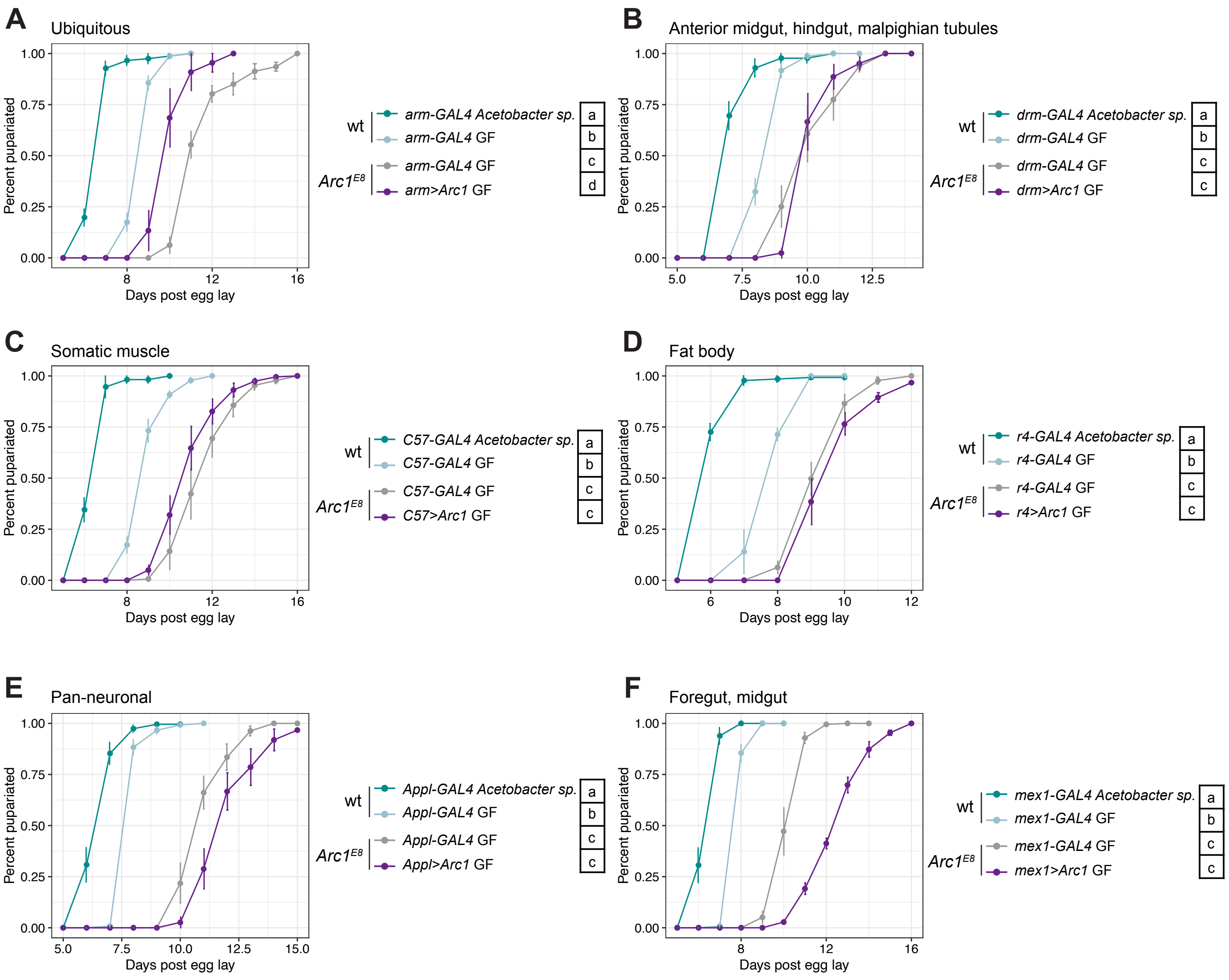


**Fig. S6. Tissue-specific Arc1 rescue under GF conditions.** Time to pupariation for GF *Arc1*$^{E8}$ animals in which Arc1 expression has been restored via the indicated tissue- and cell-type-specific GAL4 drivers. *Arc1*$^{E8}$*;GAL4* GF controls and *Arc1*$^{E8}$*;GAL4>Arc1* GF data are same as presented in Fig. 6A, but full growth curves are depicted here, along with *Acetobacter sp.*-associated and GF wild-type *GAL4* controls for reference. Control animals (*GAL4* alone in wild-type and *Arc1*$^{E8}$ backgrounds) are heterozygous for the *GAL4* transgene. Conditions sharing a letter are not statistically different from one another, one-way ANOVA with Tukey's post-hoc test.

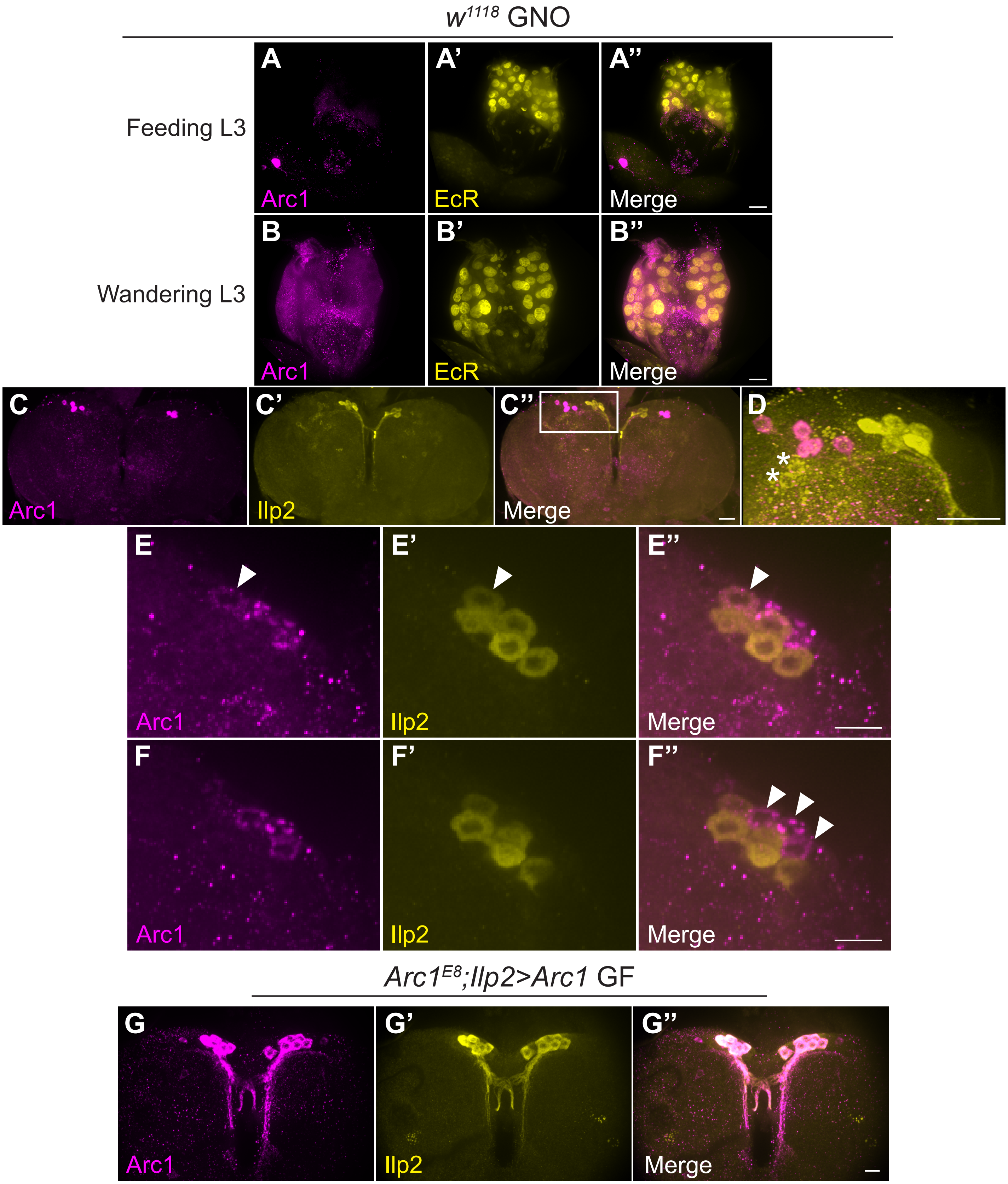


**Fig. S7. Wild-type Arc1 expression patterns in the ring gland and IPCs. A-B''.** Arc1 immunostaining level in the ring gland increases between the feeding and wandering third larval instar (L3) stages. Images were acquired with the same laser power and exposure and adjusted identically in Adobe Photoshop. Ring gland nuclei are labelled with anti-ecdysone receptor (EcR). Scale bar: 5 μm. **C-D.** Arc1-positive cells in the central larval brain shown in relation to the IPCs (labelled with anti-Ilp2). **D.** Boxed region in **C''** is shown at higher magnification in **D**. Image in **D** is from a rotated 3D projection of 125 0.2 μm slices. Two Ilp2-positive cells outside the IPC cluster and adjacent to the Arc1-positive cluster are indicated (asterisk). Scale bar: 5 μm **(C-C'')** and 5 μm **(D)**. **E-F''.** Projections of six 0.2 μm slices from two different focal planes of the same IPC cluster. Some IPCs exhibit weak Arc1 labelling (example indicated with arrowhead **E-E''**). We also observed Arc1-positive, Ilp2-negative cells adjacent to the Ilp2-positive cluster (arrowheads **F-F''**). Scale bar: 5 μm. **G-G''.** Strong, ectopic Arc1 expression in the IPCs in *Arc1$^{E8}$;Ilp2>Arc1* rescue animals (compare to Fig. 2D and S7C-F''). Scale bar: 5 μm.

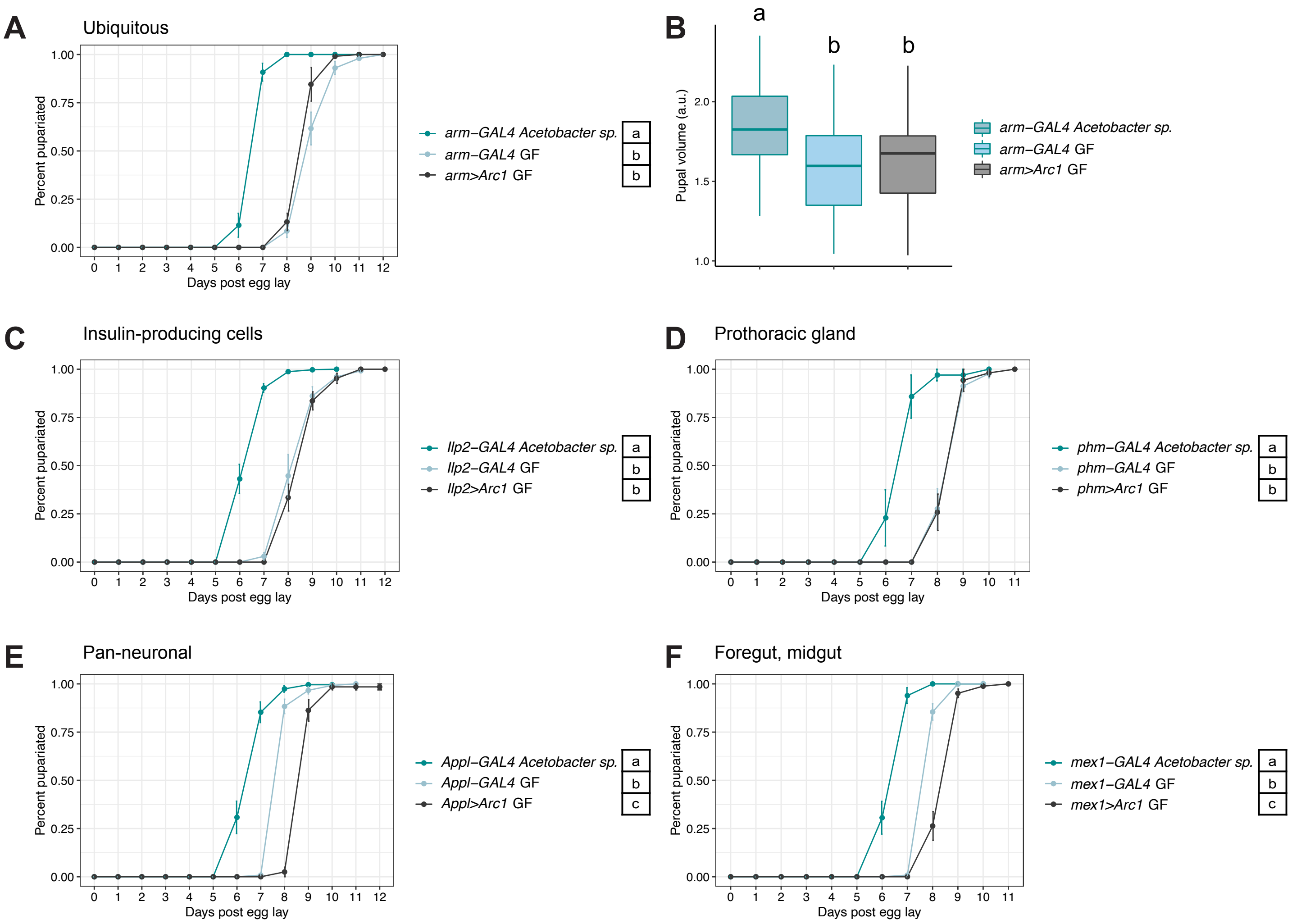


**Fig. S8. Effects of Arc1 overexpression in wild-type GF larvae on growth rate and pupal size. A, B.** Weak ubiquitous overexpression of Arc1 does not impact time to pupariation or pupal volume under GF conditions. For **B** n=37-54 pupae per condition. **C-F.** Pan-neuronal and midgut enterocyte-specific, but not IPC- or prothoracic gland-specific Arc1 overexpression delays development in GF wild-type animals. Developmental rate data for *GAL4* control animals (except GF *arm-GAL4*) are the same as in Fig. 6B,C and S6A-C, presented again here for direct comparison to overexpression animals. Conditions sharing a letter are not statistically different from one another, one-way ANOVA with Tukey's post-hoc test.

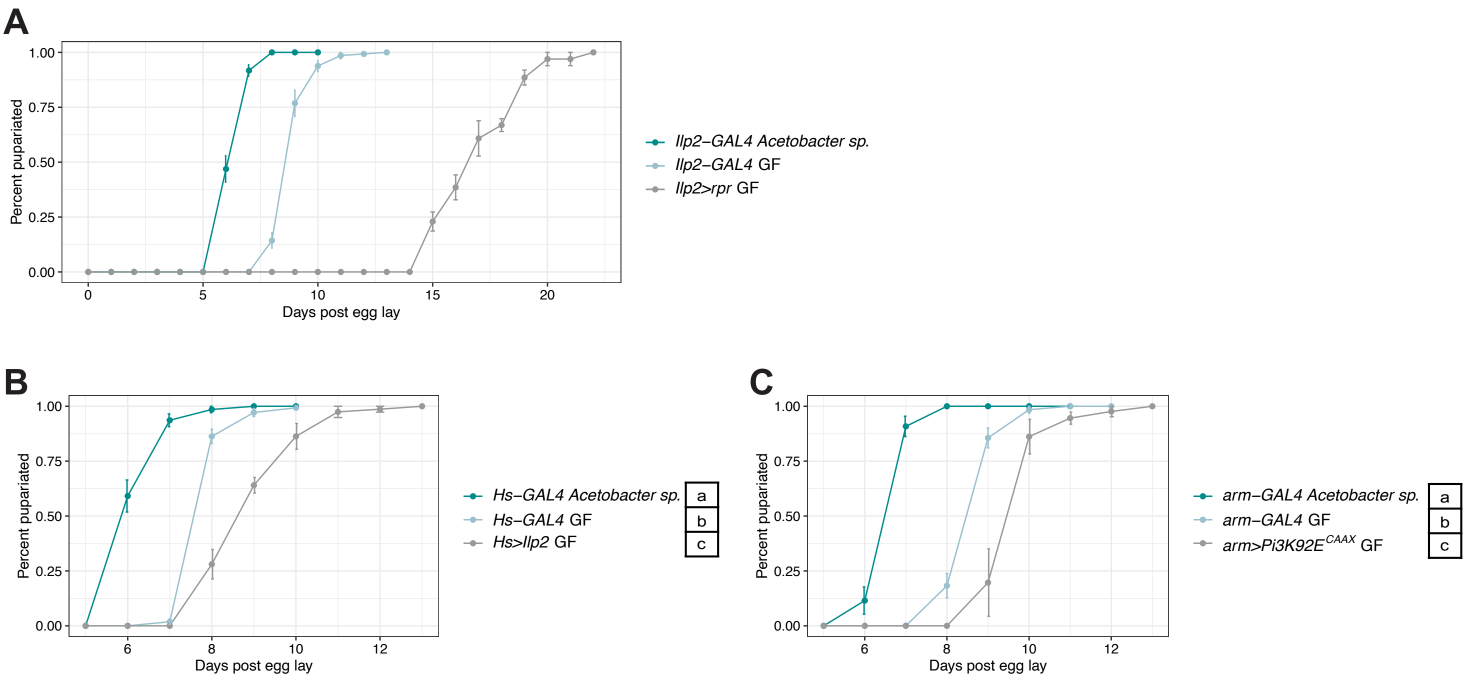


**Fig. S9. Systemic over-activation of insulin signaling or ablation of the IPCs in GF larvae induces developmental delay. A.** Expressing the pro-apoptotic *reaper* in IPCs causes severe developmental delay in GF larvae as previously shown (Rulifson et al., 2002). **B,C.** Reduced growth rate in GF larvae following IIS pathway activation via periodic heat shock-induced ubiquitous expression of Ilp2 (**B**; Brogiolo et al., 2001) or weak ubiquitous expression of membrane-targeted, constitutively active Pi3K (**C**; Homem et al., 2014). In **B** and **C**, developmental rate data for *GAL4* control animals (except GF *Ilp2-GAL4*) are same as presented in Fig. S5A and Fig. 6B, shown again here for reference. Conditions sharing a letter are not statistically different from one another, one-way ANOVA with Tukey's post-hoc test.

### Table S2. Primers used for RT-qPCR analysis

| Gene | Forward (5'-3') | Reverse (5'-3') | Reference |
|---|---|---|---|
| *Rpl32* | ATGCTAAGCTGTCGCACAAATG | GTTCGATCCGTAACCGATGT | Ponton et al., 2011 |
| *Arc1* | CATCATCGAGCACAACAACC | CTACTCCTCGTGCTGCTCCT | Mosher et al., 2015 |
| *Arc2* | CGTGGAGACGTATAAAGAGGTGG | GACCAGGTCTTGGCATCCC | FlyPrimerBank; Hu et al., 2013 |
| *Thor/4EBP* | CAGATGCCCGAGGTGTACTC | CATGAAAGCCCGCTCGTAGA | |
| *InR* | AACAGTGGCGGATTCGGTT | TACTCGGAGCATTGGAGGCAT | Obata et al., 2018 |
| *Ilp2* | GTATGGTGTGCGAGGAGTAT | TGAGTACACCCCCAAGATAG | Shin et al., 2011 |
| *Ilp3* | AAGCTCTGTGTGTATGGCTT | AGCACAATATCTCAGCACCT | |
| *Ilp5* | AGTTCTCCTGTTCCTGATCC | CAGTGAGTTCATGTGGTGAG | |
| *spok* | TATCTCTTGGGCACACTCGCTG | GCCGAGCTAAATTTCTCCGCTT | Christensen et al., 2020 |
| *sro* | CGAATCGCTGCACATGAC | TAGGCCCTGCAGCAGTTTAG | Ou et al., 2016 |
| *nvd* | GGAAGCGTTGCTGACGACTGTG | TAAAGCCGTCCACTTCCTGCGA | Pankotai et al., 2010 |
| *phm* | GGATTTCTTTCGGCGCGATGTG | TGCCTCAGTATCGAAAAGCCGT | |
| *dib* | TGCCCTCAATCCCTATCTGGTC | ACAGGGTCTTCACACCCATCTC | |
| *sad* | CCGCATTCAGCAGTCAGTGG | ACCTGCCGTGTACAAGGAGAG | |
| *shd* | CGGGCTACTCGCTTAATGCAG | AGCAGCACCACCTCCATTTC | |
